# Loss of Acid Ceramidase in Myeloid Cells Protects from Chronic Colitis in IL10-Deficent Mice

**DOI:** 10.64898/2026.01.05.697503

**Authors:** Keila S. Espinoza, Brandon K. Dahl, Jessica N. Wysong, Ella M. Ryan, Mary R. Gordon, Chelsea L. Doll, Marilyn T. Marron, Cameron A. Beard, Pujarini D. Dey, Richard J. Simpson, Justin M. Snider, Justin E. Wilson, Pawel R. Kiela, Ashley J. Snider

**Author notes:** Corresponding author: Ashley Snider 1230 N. Cherry Ave., BSRL 372, Tucson, AZ 85721.

## Abstract

**Background & Aims:** Patients with inflammatory bowel disease (IBD) exhibit elevated expression of acid ceramidase (AC), a sphingolipid metabolism enzyme. Recent studies have shown that myeloid cells contribute to the elevated expression of AC, such that the conditional loss of AC is protective in IBD.

**Methods:** Bone marrow derived macrophages (BMDMs) and neutrophils (BMDNs) were utilized to assess the role of AC in immune cell mediated inflammation. We then crossed conditional *ASAH1 LyzM^CRE^* knockout mice with *IL10* knockout mice to determine the role of AC in a model of spontaneous colitis. Colon tissues were analyzed for lipids, mRNA, and protein. We performed flow cytometry to determine the role of myeloid AC in recruiting effector T cells in disease.

**Results:** In this study, we found that loss of AC impaired secretory and migratory functions in BMDMs, but not BMDNs. Further, the conditional loss of AC protected from spontaneous, chronic colitis. Loss of AC reduced inflammatory markers, increased colon ceramides, and reduced the inflammatory metabolite sphingosine-1-phosphate (S1P). Recruitment of immune cells into intestinal tissue was significantly impaired, namely neutrophils and effector Th1/Th17 T cells.

**Conclusions:** Loss of AC reduced inflammation and impaired immune cell recruitment in chronic colitis. Targeting AC may serve as a promising therapeutic potential for patients with IBD by modulating immune cell sphingolipid metabolism.

**WHAT YOU NEED TO KNOW:** *BACKGROUND AND CONTEXT:* Acid ceramidase expression is increased in immune cells in patients with inflammatory bowel disease, specifically in macrophages.

*NEW FINDINGS:* We determined that loss of acid ceramidase (AC) in macrophages, but not neutrophils, impairs inflammatory functions *in vitro,* and that loss of AC in myeloid cells partially protects from spontaneous colitis *in vivo* by reducing immune cell recruitment into intestinal tissue.

*LIMITATIONS:* The IL10 knockout model of colitis exhibits highly variable onset and severity of disease, which may be challenging to distinguish the extent of protection.

*CLINICAL RESEARCH RELEVENCE:* This study identifies AC as a promising therapeutic target for treating inflammatory bowel disease.

*BASIC RESEARCH RELEVENCE:* This study contributes to our understanding of the role that AC plays in inflammation within immune cells, specifically myeloid cells. Additionally, this study underscores the role of immune cell sphingolipid metabolism in inflammatory bowel disease.

*LAY SUMMARY:* Loss of acid ceramidase in myeloid cells protects from chronic colitis by decreasing inflammation, altering immune cell function, and impairing the recruitment of effector immune cells to the colon.

## INTRODUCTION

Inflammatory bowel disease (IBD) is an umbrella term for persistent intestinal distress consisting of two primary derivations: Crohn’s disease (CD), which can affect the entire gastrointestinal tract, and ulcerative colitis (UC), which is restricted to the colon. Recent estimates indicate that 1.3% of adults are diagnosed with IBD^1^, which is double the prevalence of estimates since the early 2000s with a growing proportion of pediatric diagnoses^2^. While the primary cause of IBD is unknown, many factors contribute to the onset and severity of disease including diet, the gut microbiome, environmental factors, illness, and immune cell infiltration^3^. In addition, patients with IBD are 4-10x greater risk of developing colitis-associated colorectal cancer (CAC)^4, 5^, and are 1.7x higher risk of mortality^6^.

Sphingolipids are a class of bioactive metabolites ubiquitously generated in all cells with diverse roles ranging from membrane structure to cell signaling, as well as significant roles in inflammation and immune cell trafficking^7–9^. Dysregulated sphingolipid metabolism is known to influence IBD and CAC. Studies investigating the role sphingolipids in colorectal cancer (CRC) patients revealed that the expression of S1P receptors, sphingosine kinase 1^10, 11^ (SK1), and acid ceramidase^12^ (AC) were most significantly upregulated^13^. Animal studies have found that global loss of SK1 has shown to be protective in mouse models of colitis^14, 15^, whereas whole body deletion of either ceramide synthases 5 or 6 (CerS5, CesS6), as well as neutral ceramidase (nCDase), exacerbated colitis^16–18^. However, the role of sphingolipids and their metabolizing enzymes in immune cells in IBD have not been well studied. Bone marrow transplant studies have shown that SK1^19^, but not sphingomyelin synthase 2 (SMS2) ^20^ deficiency in hematopoietic cells, conferred partial protection from dextran sulfate sodium (DSS)-induced colitis. In adoptive transfer models of IBD, CerS4^21^ or CerS6^22^ deficient T cells reduced neutrophil recruitment and inflammation scoring. Further, adoptive transfer of serine palmitoyl transferase subunit 1 (SPTLC1) deficient T cells protected from colitis by impairing Th17 cell differentiation^23^. Moreover, supplementation of glucosylceramide (GluCer) laden nanoparticles rescued colitis in mice bearing T cell specific glucosylceramide synthase (GCS) deficiency^24^. This limited body of literature, namely focused on T cells, highlights the complexity of sphingolipid metabolism in IBD and our incomplete understanding the role of innate immune cell sphingolipids in IBD pathogenesis.

AC is a crucial lysosomal enzyme that breaks down ceramides into sphingosine and fatty acids, and is predominantly expressed colonic mucosal macrophages in IBD patients, and in Mac-3^+^ macrophages in murine DSS colitis^12^. The conditional loss of AC in *LyzM* expressing myeloid cells (AC^ΔMYE^) protected from DSS-induced colitis, and reduced tumor burden in azoxymethane (AOM)/DSS-induced CAC. Indeed, these studies were the first to provide insight on the role of sphingolipids in myeloid cells in IBD and CAC. In the present study, we expanded on those findings to elaborate the role of AC in a physiologically relevant model of spontaneous, chronic colitis, namely IL10 deficient (IL10^−/-^) mice. These studies distinguished a specific role of AC in macrophages but not neutrophils. Furthermore, loss of AC in myeloid cells in IL10^−/-^ mice resulted in reduced M1- and M2-like macrophage populations, and impaired the recruitment of effector T cells. These findings highlight the contribution of sphingolipid metabolism in innate immune cells and indicate that AC may serve as a promising therapeutic target for IBD.

## MATERIALS AND METHODS

### Animals

We generated AC conditional knockout mice resulting in deletion of the AC gene in the myeloid cell lineage (AC^ΔMYE^)^12^. We then crossed AC^ΔMYE^ mice with IL10 deficient mice B6.129P2-Il10^tm1Cgn^/J (Jackson Laboratory, Bar Harbor, ME) to generate AC^fl/fl^IL10^−/-^ and AC^ΔMYE^IL10^−/-^mice. Animals were maintained in a temperature-controlled environment with a 12/12hr light/dark cycle. Male and female littermates were euthanized at 8 or 24 weeks of age. Mice were monitored for body weight, rectal bleeding, and prolapse. Upon euthanasia colon tissues were flash frozen or swiss-rolled for histology. Single knockout AC^fl/fl^IL10^+/+^ and AC^ΔMYE^IL10^+/+^ mice were maintained in a specific pathogen free room with similar room conditions as described. All animal procedures were approved by the University of Arizona Institutional Animal Care and Use Committees (protocol #19-604).

### RNA Isolation and qRT-PCR

Tissues were homogenized with glass beads using MP Biomedicals FastPrep-24. RNA was isolated using the PureLink RNA isolation kit (Thermo-Fisher Scientific, Waltham, MA) and quantified via QuickDrop prior to reverse transcription utilizing qScript cDNA Synthesis Kit (Quantabio, Beverly, MA) at 1 µg/µl. cDNA was used for quantitative real-time RTPCR performed in triplicate with FAM probe primers (Thermo-Fisher Scientific). Expression levels of genes of interest were normalized to VIC probe β-actin mRNA.

### Protein Isolation and ELISAs

Tissues were homogenized with glass beads using MP Biomedicals FastPrep-24 in 400µl of lysis buffer containing cOmplete™ EDTA-free protease inhibitor (Millipore Sigma, Carlsbad, CA). Supernatant and tissue were transferred into Eppendorf tubes and beads were rinsed with an additional 400µl of buffer. Samples were sonicated and centrifuged at 15000 x g for 15 min at 4 °C. Supernatant was collected and stored in aliquots to avoid repeated freeze-thaw cycles. Protein concentration was quantified by Pierce BCA assay. ELISA buffer and antibody kits (Thermo-Fisher Scientific) were acquired and performed on tissue lysate or cell culture media according to manufacturer instructions.

### Histology

Swiss-rolled colons were incubated in buffered formalin overnight prior to transfer into 70% ethanol. Tissue embedding, slide mounting, and H&E staining were performed by the University of Arizona Cancer Center (UACC) Tissue Acquisition and Cellular/Molecular Analysis Shared Resource (TACMASR). H&E stained slides were analyzed by veterinary pathologist Dr. David Besselsen, blinded to the study design and groups. For DAB staining, slides were deparaffinized and serially rehydrated. Antigen retrieval was performed at 110°C in sodium citrate buffer for 10min. Slides were incubated in horseradish peroxidase for 15 minutes, then in 10% goat serum to block nonspecific binding. Slides were incubated with primary antibody overnight at 4 °C. HRP linking was performed using the Rabbit HRP/DAB kit (Abcam, Waltham, MA) followed by DAB substrate development (Thermo-Fisher Scientific) according to manufacturer instructions. Slides were briefly counterstained in Mayer’s hematoxylin prior to serial dehydration and mounting. Slides were imaged at 20x and analyzed using Fiji software by a researcher blinded to the experimental groups.

### FITC-Dextran Permeability Assay

4 hrs prior to euthanasia, mice were administered 600 mg/kg FITC-dextran (Millipore Sigma) via oral gavage. Blood was collected, allowed to coagulate, then centrifuged at 2000 xg for 12 min. Serum was collected and diluted 1:5 in PBS. 100 µl of sample was plated in an opaque 96-well plate and read on a fluorescent plate reader (excitation 485 nm, emission 535 nm).

### Lipid Mass Spectrometry

Lipids from tissues and cell culture experiments were extracted as previously described^25^. Analyses were conducted by the UACC Analytical Chemistry Shared Resource Core. Data were normalized to total lipid phosphate (Pi) present in the organic phase^26^ of the Bligh and Dyer extraction^27^ detected by phosphomolybdate assay. Sphingolipid levels are expressed as pmol/nmol of lipid Pi.

### Flow Cytometry

Upon euthanasia, blood was collected into EDTA tubes to prevent coagulation. Blood samples were treated twice with red blood cell (RBC) lysis buffer (Fisher Scientific, Hampton, NH). Spleens were excised and mashed on a petri dish over a 70 µm filter. Spleen samples were treated once with RBC lysis buffer after centrifugation. Colon tissues were collected and processed according to the published protocol^28^. For surface staining (CD45, CD3, CD4, CD8, CD19, CD11b, CD11c, MHC II, F480, Ly6G, CD25, Ly6C, CD206), samples were pelleted in a 96-well round bottom plate and resuspended in FC block for 10 min. Antibody cocktail was incubated at 4 ° for 1 hr. Samples were rinsed and fixed for 10 min in formalin at RT, then transferred to FACS tubes. For intracellular staining (FoxP3, IFNg, IL-17A, Granzyme B), samples first incubated in 4 µM of Monensin (Thermo-Fisher Scientific) for 3 hr in complete RPMI media at 37 °C. After surface staining, samples fixed for 30 min in formalin at room temperature, then rinsed in 1 x permeabilization buffer (Thermo-Fisher Scientific). Intracellular antibodies were added at 1:250 and incubated overnight at 4 °C. Samples were rinsed in permeabilization buffer and transferred to FACS tubes. All samples were read on a BD LSR Fortessa provided by the University of Arizona Department of Pediatrics. Data were exported and analyzed using FlowJo v10.8 software (BD Life Sciences, Franklin Lakes, NJ).

### Isolation and Culture of Bone Marrow-Derived Macrophages (BMDMs) and Neutrophils (BMDNs)

Bone marrow was isolated from 10-week-old AC^fl/fl^ and AC^_MYE^ mice by flushing both femurs and tibias with room temperature complete RPMI containing 2 mM EDTA. Female mice were used for BMDMs. Male mice were used for BMDNs. For BMDM generation, bone marrow was briefly treated with RBC lysis buffer. 3.0 ×10^6^ cells were plated in a 10 cm petri dish containing 9 mL DMEM + 20 ng/mL M-CSF. On day three of culture, 5 mL of additional M-CSF media was replenished. Cells were lifted on day six of culture by incubating in chilled PBS + 5 mM EDTA at 4 °C for 10 min. Macrophages exhibited over 90% viability and 95% purity by FACS analysis. To obtain CD11b^+^Ly6G^+^-enriched population of BMDNs, we utilized EasySep™ Mouse Neutrophil Enrichment Kit (StemCell, Cambridge, MA) according to the manufacture instructions. Neutrophils exhibited over 95% viability and 90% purity by FACS analysis.

### LPS Assays

To assess inflammatory response, 0.8×10^6^-1.0×10^6^ cells were plated in 6 well plates – BMDNs remained undisturbed for at least 3 hr in 2 mL of complete RMPI, whereas BMDMs adhered overnight in complete DMEM. For BMDNs, 1.5 mL of RPMI + LPS (Millipore Sigma) was added to a final concentration of 25 ng/mL for indicated times. Cells were resuspended in media and collected in a 15 mL tube. BMDNs were pelleted and media was transferred to be stored at -20 °C. Cells were then resuspended in 2 mL of cell extraction solvent (2:3 70% isopropanol:EtOAc) after removal of media. For BMDMs, media was aspirated and 3 mL of DMEM + 25 ng/mL LPS was added. Media was aliquoted and stored at -20°C. Cells were rinsed in ammonium formate and scraped with 2 mL cell extraction solvent prior to being stored at -20 °C.

### Migration Assay

Migration assay was performed according to the published protocol^29^. C5a and MCP1 (Thermo-Fisher Scientific) were utilized as chemoattractants at 85 ng/mL and 125 ng/mL, respectively. BMDNs migrated for 10 hr. BMDMs migrated for 6 hr.

### Phagocytosis Assay

Phagocytosis assay kit (Cayman Chemical, Ann Arbor, MI) was performed according to the manufacture suggestions. For BMDNs, cells cultured for 4 hr in complete RPMI containing latex beads-rabbit IgG-FITC complex at 1:200 dilution. For BMDMs, cells cultured for 4 hr with beads at 1:500 dilution. Cells were then rinsed, fixed, and assessed by FACS analysis.

### BMDM Polarization

BMDMs were polarized for 16 hr using 8 mL complete DMEM containing 10 ng/mL LPS + 50 ng/mL IFNg for M1 phenotype, or 10 ng/mL IL-4 + 10 ng/mL IL-13 for M2 phenotype. Polarized cells were functionally assessed as described above.

### Statistical Analysis

Analysis Software GraphPad Prism v10.6 (Boston, MA) was used for statistical analysis and graph generation. Data were analyzed using Two-Way ANOVA followed by Fisher’s LSD multiple comparison post-hoc test when multiple groups were compared. An unpaired T-test was used when only comparing two groups. Outliers were identified using the Grubbs’ outlier test with an alpha of 5%.

Schematics generated via BioRender (San Francisco, CA).

This study adheres to the ARRIVE 2.0 guidelines.

## RESULTS

### AC regulates inflammation in macrophages, but not neutrophils

Both neutrophils and macrophages express *LyzM* and contribute greatly to persistent inflammation during colitis. To determine the extent that AC differentially regulates sphingolipid metabolism (Figure 1A) and inflammatory responses in these immune cells, we isolated neutrophils (BMDNs) and generated macrophages (BMDMs) from the bone marrow of AC^fl/fl^ and AC^ΔMYE^ mice. We first assessed how sphingolipid metabolism changed before and after eliciting an inflammatory response by stimulating cells with LPS. Both AC^ΔMYE^ myeloid cell types exhibited elevated dihydroceramides and ceramides (Figure 1B and D) as compared to AC^fl/fl^ and exhibited increased ceramide generation upon LPS stimulation. Sphingosine and S1P were unaltered in AC^ΔMYE^ BMDNs (Figure 1C), whereas BMDMs exhibited reduced S1P generation (Figure 1E). Notably, ceramide species in AC^ΔMYE^ BMDNs were elevated ∼5-fold as compared to AC^fl/fl^ (Figure 1F). Increased dihydroceramides and ceramides were even further elevated in AC^ΔMYE^ BMDMs (some species by 20-fold) (Figure 1G). These data indicate that AC differentially regulates sphingolipid metabolism dependent on cell type.

**Figure 1.**
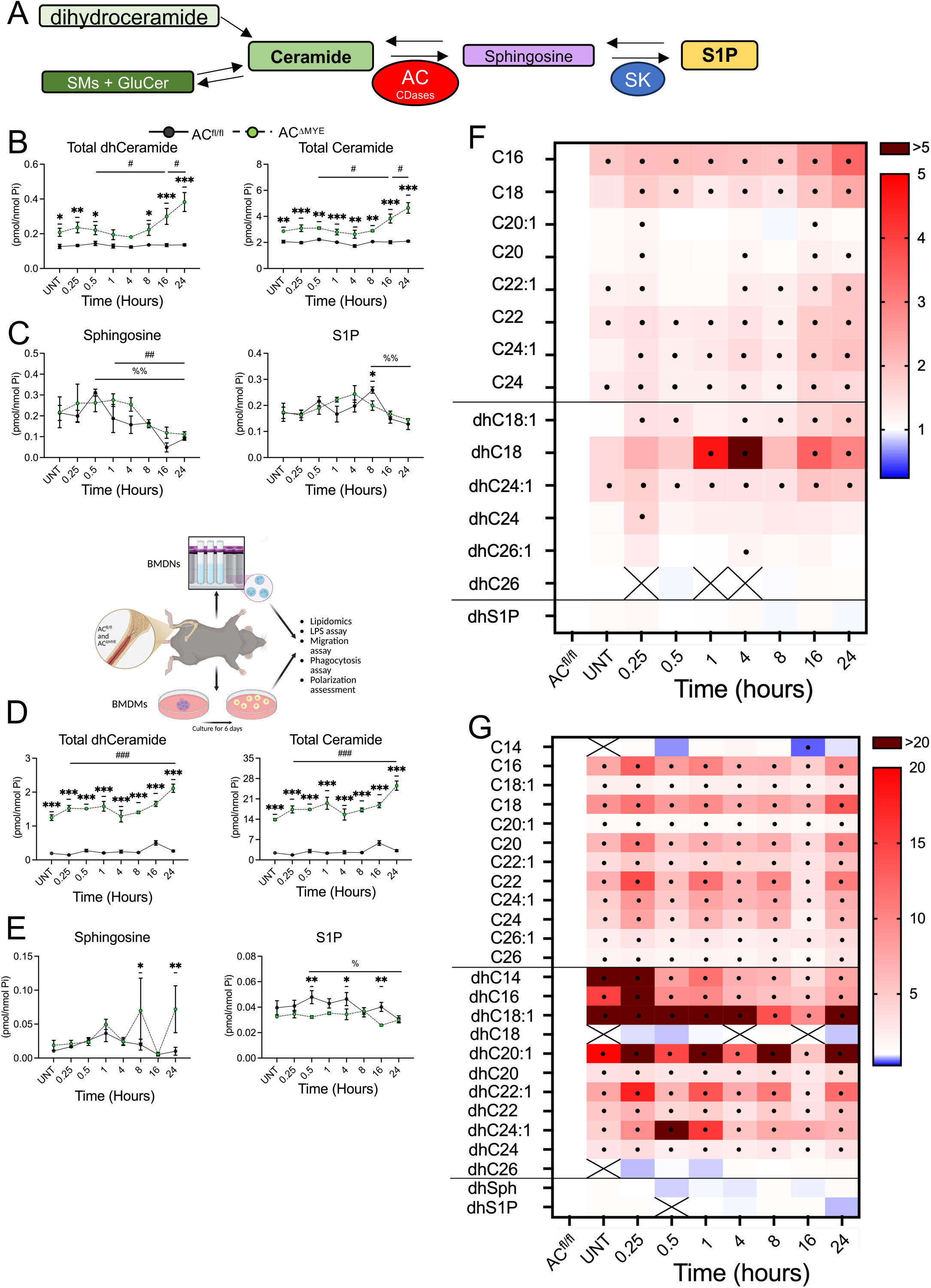
Loss of AC reduces S1P and increases ceramide generation in myeloid cells. AC^fl/fl^ and AC^DMYE^ BMDNs and BMDMs were stimulated with 25ng/mL LPS for the indicated times. (A) Simplified scheme for sphingolipid metabolism. BMDN total (B) dihydroceramide, ceramides, and (C) sphingoid bases, n≥3. BMDM total (D) dihydroceramide, ceramides, and (E) sphingoid bases, n=3. Data represent mean ± SEM; *p<0.05, **p<0.01, ***p<0.001 between genotype; #p<0.05, ##p<0.01, ###p<0.001 within AC^DMYE^; %p<0.05, %%p<0.01, %%%p<0.001 within AC^fl/fl^. Specific species at each time point for AC^DMYE^ (F) BMDNs and (G) BMDMs normalized to AC^fl/fl^. 2-way analysis of variance (ANOVA), Šídák multiple comparisons correction; • indicates p<0.05. X indicates insufficient abundance.

Ceramides and S1P are bioactive signaling lipids which regulate cell signaling and inflammation. Given that loss of AC significantly elevated ceramides, we sought to interrogate the functional role of AC in BMDNs and BMDMs. Loss of AC did not significantly alter migration, phagocytosis, or cytokine secretion in BMDNs (Supplementary Figure 1). However, loss of AC in BMDMs resulted in significant impairment of migration towards C5a (Figure 2A) and reduced percentage macrophages carrying out phagocytosis, though the number of particles (engulfed beads) per macrophage were similar between genotypes (Figure 2B). Cytokine and chemokine secretion were also significantly ablated in AC^ΔMYE^ BMDMs up to 24 hours after LPS stimulation (Figure 2C). Indeed, these data suggest that while AC may increase ceramide levels in neutrophils, AC selectively regulates inflammatory responses in macrophages.

**Figure 2.**
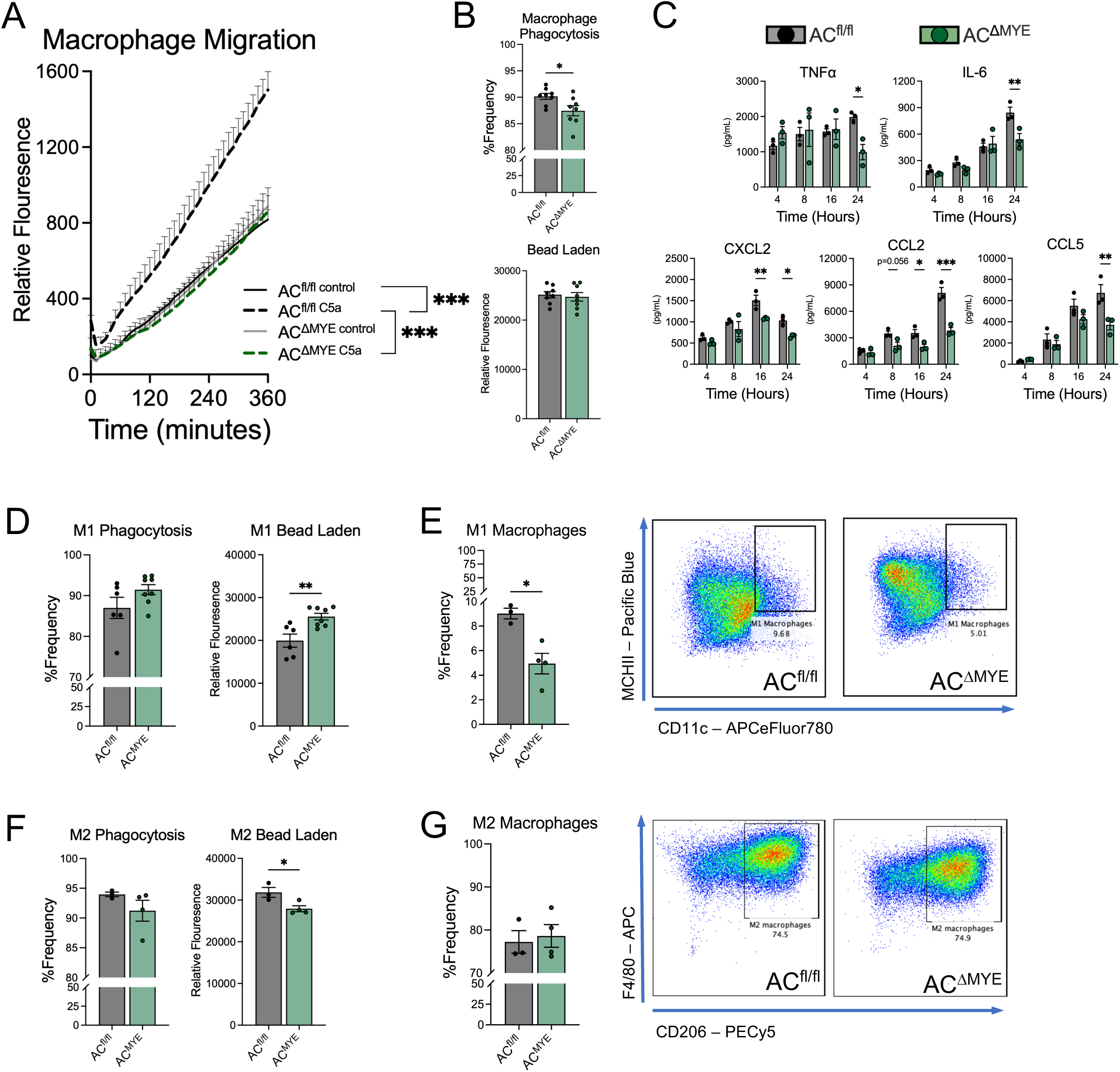
AC selectively regulates inflammatory response in macrophages, but not neutrophils. (A) BMDM migration assay against C5a, terminal timepoint assessed by 2-way ANOVA. (B) BMDM phagocytosis assay, unpaired t test: percent phagocytic cells (top) and relative particle engulfment (bottom). (C) Post-LPS stimulation cytokine and chemokine media assessment via ELISA, 2-way ANOVA, Šídák multiple comparisons correction. (D) M1 polarized BMDM phagocytosis assay, unpaired t test: percent phagocytic cells (left) and relative particle engulfment (right). (E) MHCII+ CD11c+ Type M1 BMDM polarization frequency phenotyping, unpaired t test. (F) M2 polarized BMDM phagocytosis assay, unpaired t test. (G) F4/80+ CD206+ Type M2 BMDM polarization frequency phenotyping, unpaired t test. Data represent mean ± SEM, n≥3; *p<0.05, **p<0.01, ***p<0.001.

We then sought to further define the role of AC in macrophages polarized *in vitro* towards M1 or M2 phenotypes. Interestingly, migration ability of either M1- or M2-polarized macrophages was not dependent on AC expression (Supplementary Figure 2A and B). While the phagocytic ability in M1 polarized macrophages did not differ between genotypes, AC^ΔMYE^ M1 macrophages exhibited significantly higher phagocytosis (bead engulfment) (Figure. 2D). AC^ΔMYE^ BMDMs were resistant to M1-polarization induced by LPS and IFNg, as demonstrated by reduced frequency of MHCII^high^CD11c^+^ macrophages (Figure. 2E). Conversely, AC^ΔMYE^ BMDMs polarized towards M2 phenotype with IL-4 and IL-13 showed significantly lower phagocytic ability as compared to AC^fl/fl^ controls (Figure. 2F), while M2 polarization frequency defined by CD206 cell surface expression was not different between genotypes (Fig. 2G). These data indicate that AC plays a major role in promoting the inflammatory response in macrophages prior to and after M1/M2 polarization.

### Loss of acid ceramidase in myeloid cells protects from colitis in IL10^−/-^ mice

To advance the pathophysiologic relevance of myeloid derived AC as a key driver of chronic colitis, we utilized IL10^−/-^ mice which spontaneously develop colitis, thereby modeling the chronic nature of human disease. Body weight, intestinal permeability, and histology were not significantly different between AC^fl/fl^IL10^−/-^ and AC^ΔMYE^IL10^−/-^ mice (Figure 3A; Supplementary Figure 3). However, AC^fl/fl^IL10^−/-^ mice demonstrated significantly decreased body weight at 24 weeks as compared to AC^fl/fl^IL10^+/+^ mice, while final body weights were not different between AC^ΔMYE^IL10^−/-^ and AC^ΔMYE^IL10^+/+^ mice, suggesting partial protection from weight loss in AC^ΔMYE^IL10^−/-^ mice. Similarly, AC^ΔMYE^IL10^−/-^ mice were protected from splenomegaly and colon shortening (Figure 3C and D). Proinflammatory markers in colon tissues significantly increased in expression between 8 and 24 weeks in AC^fl/fl^IL10^−/-^ mice (Figure 3E & F); however, this was ablated in AC^ΔMYE^IL10^−/-^ mice. Furthermore, the expression of pSTAT3, which has been implicated downstream of S1P, was elevated in AC^fl/fl^IL10^−/-^ mice, but not in AC^ΔMYE^IL10^−/-^ mice (Figure 1G). Lastly, we observed sexual dimorphism suggesting that ablating AC expression may be more protective in male myeloid cells than in females (Supplementary Figure 4). Collectively, these data strongly suggest that loss of AC in myeloid cells restricts chronic colitis.

**Figure 3.**
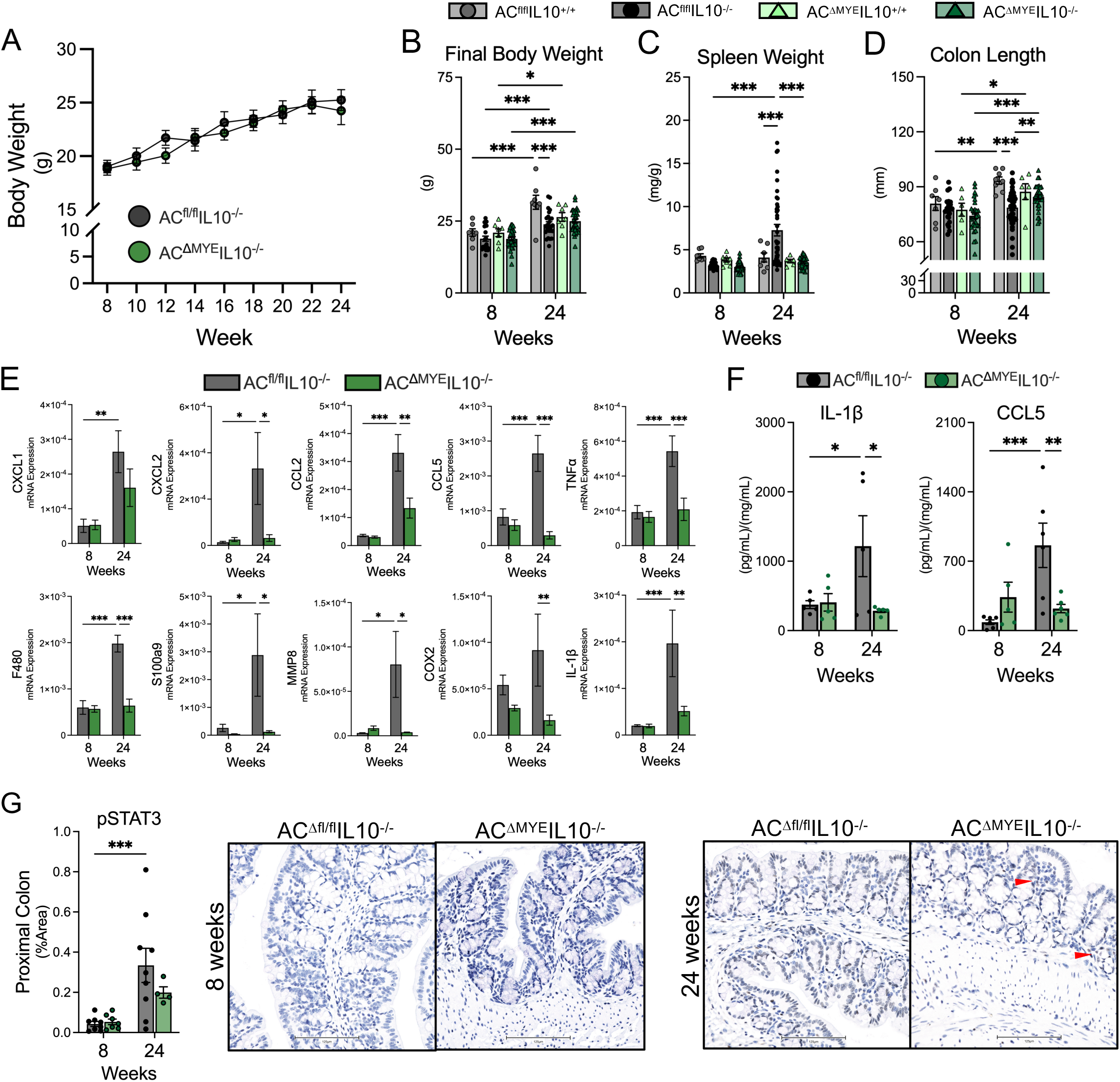
Loss of acid ceramidase in myeloid cells protects from inflammation and parameters of colitis. Male and female AC^fl/fl^IL10^−/-^ and AC^ΔMYE^IL10^−/-^ mice were aged to 8 and 24 weeks of age. (A) Body weight over time, n≥11. (B) Terminal body weight, (C) spleen weight normalized to body weight, and (D) colon length, n≥22; AC^fl/fl^IL10^+/+^ and AC^ΔMYE^IL10^+/+^ mice were aged to corresponding timepoints for disease onset comparisons, n≥7. (E) rt-PCR mRNA analysis of colon tissue, n≥7. (F) ELISA analysis of colon tissue lysate, n≥5. (G) Histological assessment of proximal colon tissue expression of p-STAT3, n≥4. 2-way ANOVA, Fisher’s LSD test. A minimum of 3 mice were used per sex per genotype per time point for all experimental endpoints. Data represent mean ± SEM; *p<0.05, **p<0.01, ***p<0.001.

### Myeloid AC regulates sphingolipid levels in colon tissues

As AC degrades ceramide and can provide a substrate for the generation of S1P, we assessed sphingolipids in the colon. Ceramides were elevated in AC^ΔMYE^IL10^−/-^ mice at 8 and 24 weeks of age (Figure 4A), although this did not reach significance at the 24 week timepoint. Assessment of specific chain length ceramides revealed significant increases in C16 and C18 ceramides at 8 weeks, and C16 ceramide remained elevated in AC^ΔMYE^IL10^−/-^ mice at 24 weeks (Figure 4B). Sphinganine and sphingosine, sphingoid bases and products of AC, were reduced in AC^ΔMYE^IL10^−/-^ mice, as were dhS1P and S1P at either 8 or 24 weeks (Figure 4C and 4D). Dihydroceramides nor glucoslyceramides were altered by the loss of AC (Supplementary Figure 5). Sphingomyelins (SMs) exhibited an age dependent reduction in both genotypes, though SMs were elevated in AC^ΔMYE^IL10^−/-^ mice as compared to AC^fl/fl^IL10^−/-^ mice at 24 weeks (Figure 4E). Examination of SM species revealed significant increases in C18 and C20 SMs at 24 weeks (Figure 4F). Interestingly, lyso-SM, which can be generated by AC, was reduced in AC^ΔMYE^IL10^−/-^mice in chronic disease (Figure 4G). These data emphasize the contribution of sphingolipid metabolism specifically in myeloid cells to colonic sphingolipid metabolism.

**Figure 4.**
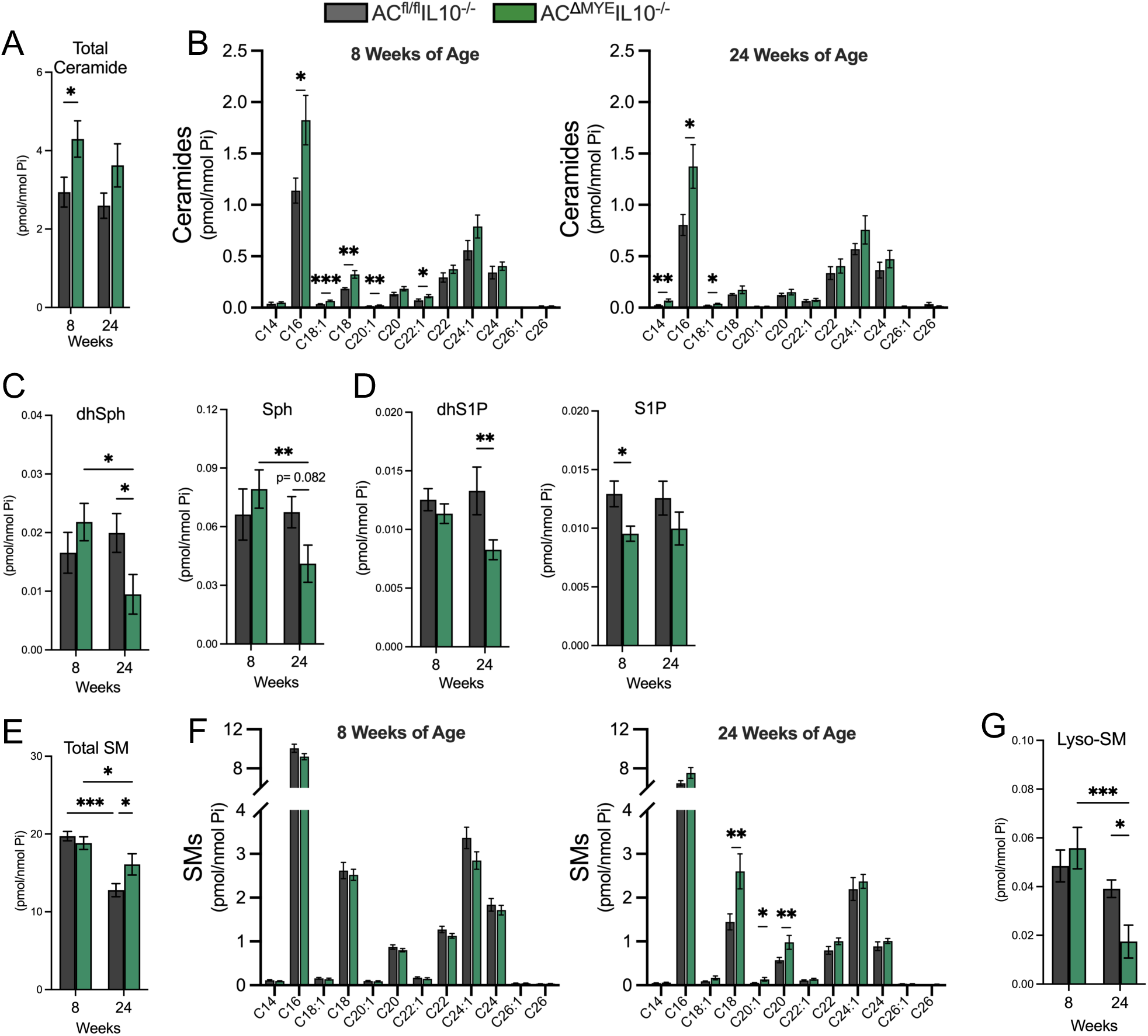
Loss of AC increases protective sphingolipids and reduces generation of S1P in colon tissue. Colon tissue lipids of AC^fl/fl^IL10^−/-^ and AC^ΔMYE^IL10^−/-^ mice were assessed at 8 and 24 weeks of age normalized to Pi. (A) Total ceramides and (B) species distribution at 8 or 24 weeks of age. (C) dhSph and Sph. (D) dhS1P and S1P. (E) Total SM and (F) species distribution at 8 or 24 weeks of age. (G) Lyso-SM. and 2-way ANOVA, Fisher’s LSD test. Data represent mean ± SEM, n≥8; *p<0.05, **p<0.01, ***p<0.001.

### Myeloid AC regulates both innate and adaptive immune cell recruitment to colon tissue

Our BMDM data indicated that loss of AC significantly impairs secretion of inflammatory markers and migratory abilities in macrophages, which may extend to the recruitment of other immune cells to sites of inflammation. To determine the role of myeloid cell AC on immune cell recruitment during colitis, we performed flow cytometry on blood, spleen and colon tissues in AC^fl/fl^IL10^−/-^ and AC^ΔMYE^IL10^−/-^ mice (Supplementary Figure 6). Loss of myeloid cell AC exerted minimal impacts on T and B lymphocytes (Supplementary Figure 7). Splenic macrophages (Figure 5A) and blood monocytes (Figure 5B) were significantly reduced in AC^ΔMYE^IL10^−/-^ mice, but macrophages in the colon were not altered between genotypes (Figure 5C and D). Interestingly, although neutrophils were reduced in the spleen (Figure 5E), circulating blood neutrophils were significantly elevated in AC^ΔMYE^IL10^−/-^ mice (Figure 5F). Comparatively, neutrophil infiltration in the colonic lamina propria (LP) and intra-epithelial compartments (IEC) were significantly reduced in the colons of AC^ΔMYE^IL10^−/-^ mice at 24 weeks (Figure 5G & H). These data suggest that while neutrophils may be elevated in circulation, loss of AC in myeloid cells may restrict their recruitment into colonic mucosa, likely due to reduced chemokine expression by AC-deficient macrophages (Figure. 2C).

**Figure 5.**
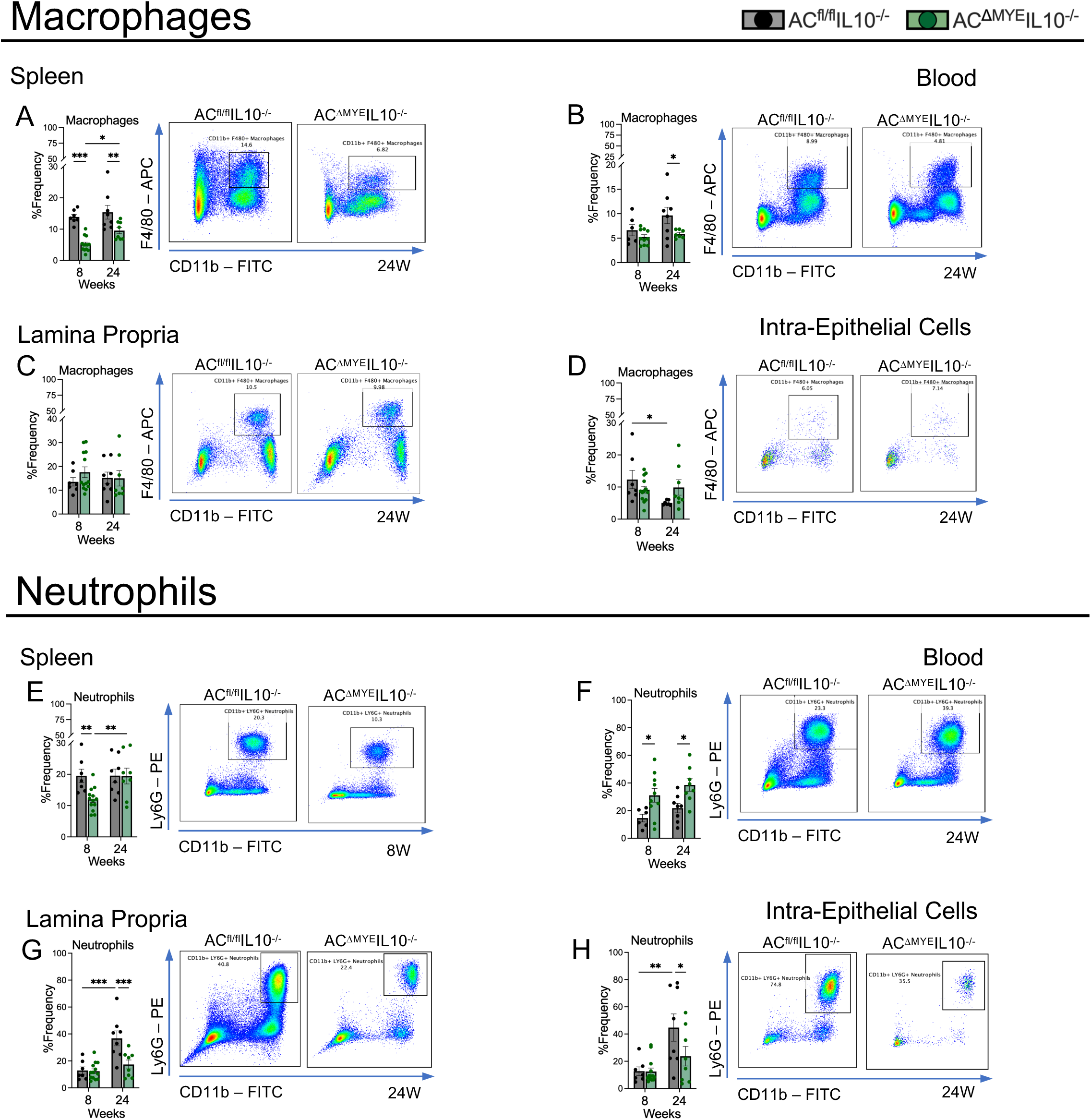
AC regulates macrophage and neutrophil populations in chronic colitis. Spleen, blood, and colon tissue compartments of the lamina propria (LP) and intra-epithelial cell (IEC) layers of AC^fl/fl^IL10^−/-^ and AC^ΔMYE^IL10^−/-^ mice were assessed by flow cytometry at 8 and 24 weeks of age. F4/80+CD11b+ macrophages in the (A) spleen, (B) blood, colon (C) LP and (D) IEC tissues. Ly6G+CD11b+ neutrophils in (E) spleen and (F) blood, colon (G) LP and (H) IEC tissues. Representative dot plots to the right of quantified graphs. 2-way ANOVA, Fisher’s LSD test. Data represent mean ± SEM, n≥6; *p<0.05, **p<0.01, ***p<0.001.

As innate immune cells, macrophages are the first line of defense against inflammation and pathogens, and can polarize further into a spectrum of effector macrophages, simplified as M1-or M2-like cells. Additionally, macrophages inform the adaptive immune branch to mount stronger responses mediated by effector T cell subsets. To determine the role of AC in effector cell recruitment during colitis, we then assessed proinflammatory M1-like and proresolving M2-like macrophages, Th1, and Th17 T cells in blood and colon tissues of AC^fl/fl^IL10^−/-^ and AC^ΔMYE^IL10^−/-^ mice (Supplementary Figure 8). M1- and M2-like macrophages were predominantly reduced in AC^ΔMYE^IL10^−/-^ mice at 8 weeks of age in spleen, blood and colon tissues (Figure 6A-D). Assessment of effector T cells identified that although overall CD4+CD25+ T cell populations were unaltered, CD8+GranzymeB+ cytotoxic T cells were reduced throughout all tissues at varying time points in AC^ΔMYE^IL10^−/-^ mice (Supplementary Figure 9). Furthermore, while regulatory FOXP3+ T cells (T_regs_) remained unchanged between genotypes, IFNg+T were predominantly reduced in blood and spleen of AC^ΔMYE^IL10^−/-^ mice, while IL-17A+T were reduced in the colonic LP of AC^ΔMYE^IL10^−/-^ mice. Interestingly, Th1 and Th17 T cells were reduced in AC^ΔMYE^IL10^−/-^ mice in chronic colitis across all studied tissues (Figure 6E-H). These data indicate that myeloid cell AC may play a role in regulating T cell recruitment and/or differentiation towards Th1/Th17 phenotypes.

**Figure 6.**
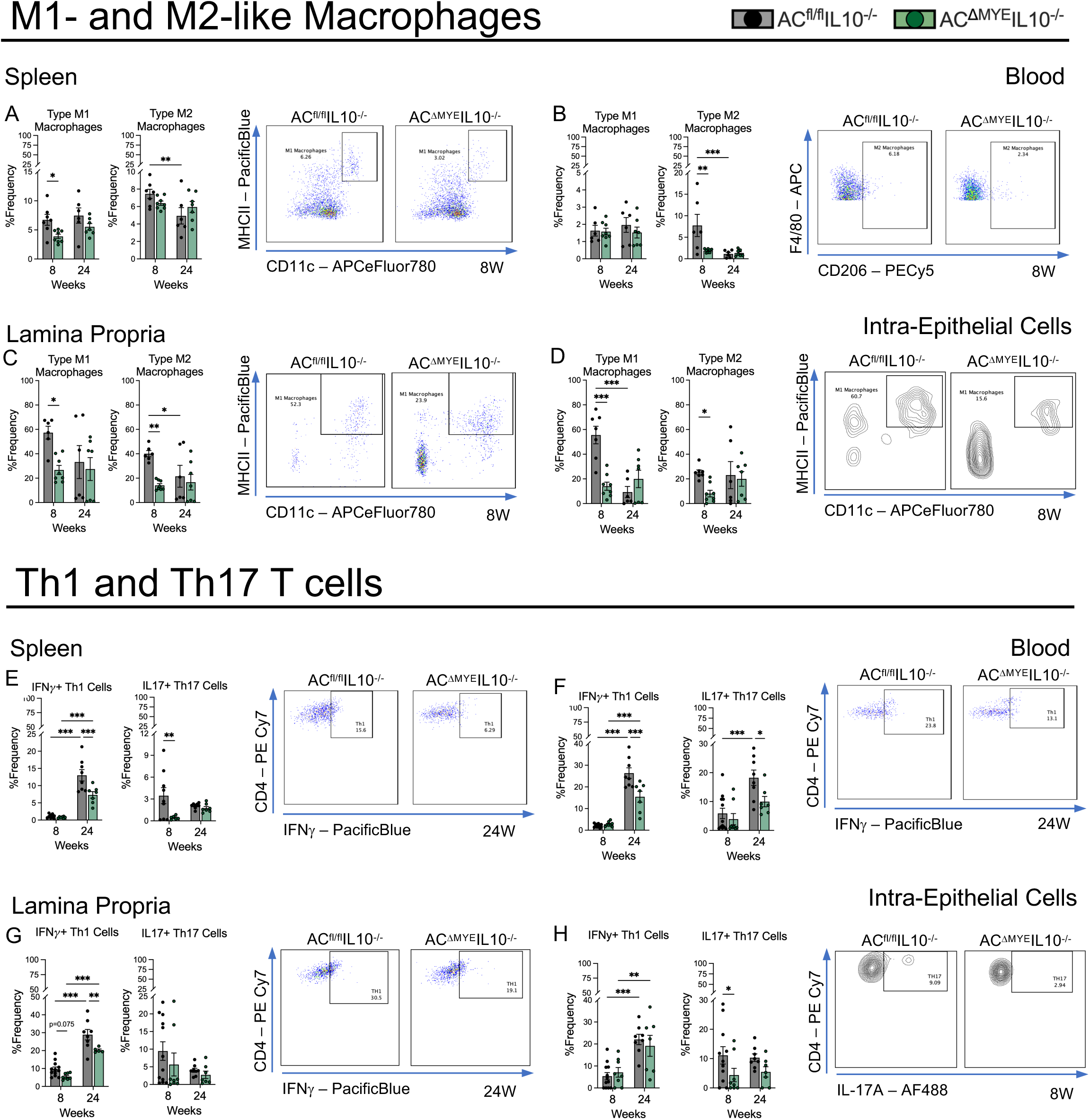
AC regulates recruitment of effector myeloid and T cells throughout tissues. F4/80+CD11b+ MHCII+CD11c+ Type M1 macrophages and F4/80+CD11b+ CD206+ Type M2 macrophages in (A) spleen, (B) blood, colon (C) LP and (D) IEC tissues, n≥6. CD4+CD25+ IFNg+ Th1 and CD4+CD25+ IL-17A+ Th17 T cells in (E) spleen, (F) blood, colon (G) LP and (H) IEC tissues, n≥7. Representative dot plots to the right of quantified graphs. 2-way ANOVA, Fisher’s LSD test. Data represent mean ± SEM; *p<0.05, **p<0.01, ***p<0.001.

## DISCUSSION

The impact of sphingolipid metabolism in IBD and CAC has been extensively studied with the use of genetically engineered mouse models and inhibitors (reviewed in ^30^). However, the specific role of immuno-sphingolipid metabolism in IBD remains elusive. In this study we set out to define the role of AC in myeloid cells in inflammation and chronic colitis in IL10^−/-^ mice. Our data demonstrate that loss of AC in BMDMs reduced secretion of pro-inflammatory cytokines and chemokines, blunted migration and phagocytosis, and impaired type M1 polarization. Loss of AC in myeloid cells *in vivo* reduced the severity of colitis, increased ceramide levels in colonic tissue, and reduced recruitment of immune cells to the colon. This study also highlights the differential role of AC in macrophages and neutrophils *in vitro*, emphasizing the role of AC regulating inflammatory pathways specifically in macrophages. Further, our findings from an *in vivo* model of chronic colitis show that loss of AC in macrophages may reshape the intestinal environment to reduce inflammation, as well as decrease neutrophil and T cell infiltration (Figure 7).

**Figure 7.**
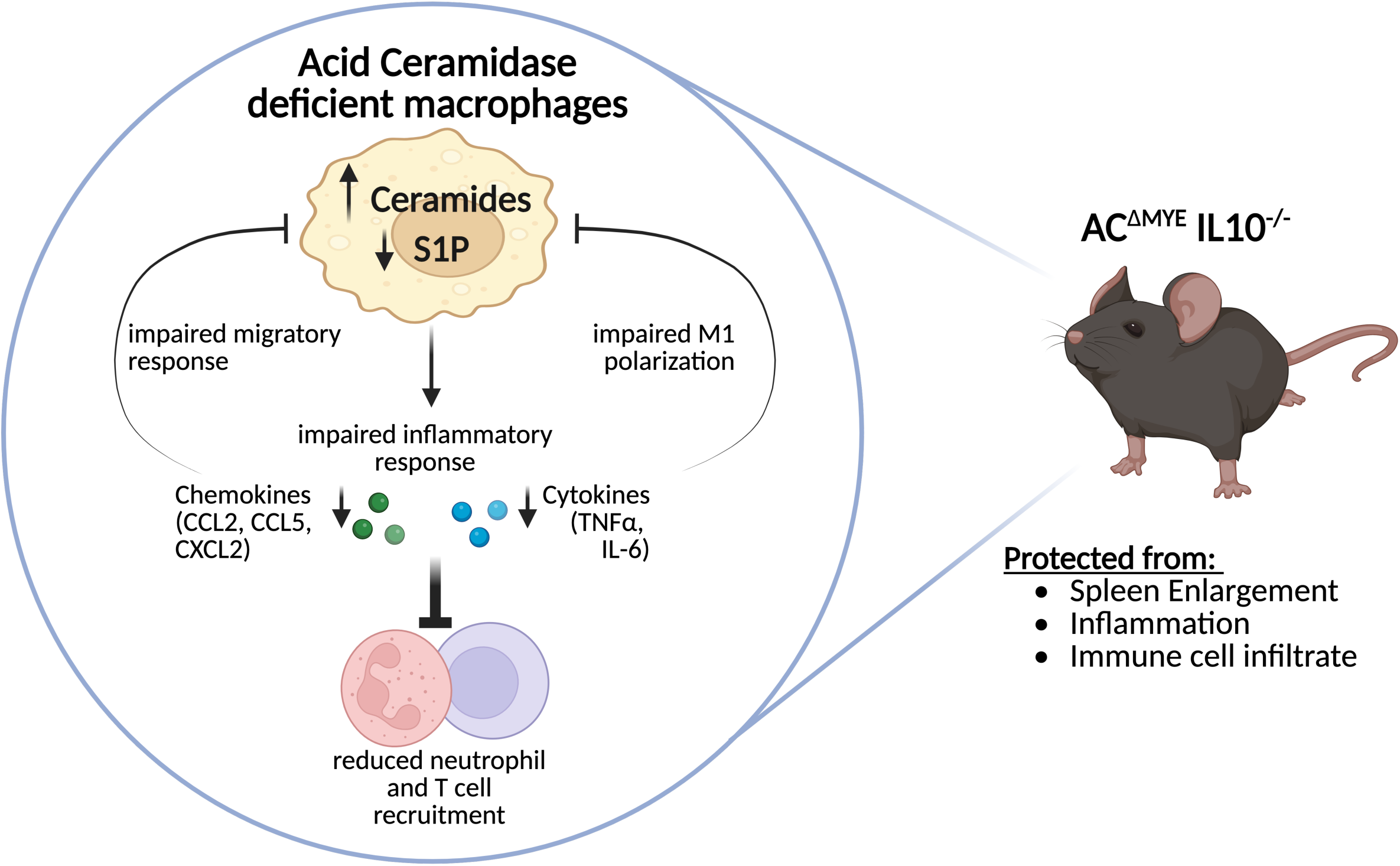
Proposed mechanism on the loss of AC in macrophages downregulating inflammatory responses to attenuate chronic colitis.

Our previous work demonstrated that loss of AC in myeloid cells reduced neutrophil recruitment in an acute, chemically-driven model of colitis^31^; however, primary BMDNs did not exhibit reduced migration in response to stimuli^31^. In this study we expanded these findings to examine neutrophil function and cytokine secretion, both of which were unchanged with loss of AC.

Though we are the first to report the role of AC in neutrophils to this detail, AC has been shown to regulate sphingolipid levels and immune responses in macrophages^12, 32–34^. Indeed, in our study BMDMs lacking AC exhibited significant increases in ceramide levels ^12^. Loss of neutral ceramidase (nCDase) in macrophages resulted in reduced sphingosine levels^35^. Interestingly, loss of SKs similarly elevated ceramides and ablated S1P, but did not change BMDM function^36^. We previously demonstrated that loss of AC in BMDMs reduced TNFα and IL-6 secretion in response to LPS^12^, these studies extended those findings to show that loss of AC also reduces generation of chemokines (CXCL2, CCL2 and CCL5). Indeed, our work in breast cancer has also implicated AC in the regulation of CCL5^37^. Secretion of IL1-β was also impaired nCDase deficient macrophages in response to infection^35^. Further, these macrophages exhibited poor anti-tumor response against breast cancer by promoting CD8+ cytotoxic T cell exhausion^38^. In contrast, myeloid cells lacking alkaline ceramidase (*ACER3*) demonstrated enhanced expression of inflammatory markers *in vitro*^39^, identifying variable roles for ceramidases in myeloid-derived cells. Together these studies suggest that sphingolipid enzymes may regulate immune cell function partially independent of sphingolipid levels.

Our data suggest that AC may regulate the response of macrophages towards polarizing cytokines. Ceramides have been shown to be crucial in polarizing macrophages towards a more aggressive M1 phenotype^40–42^. Previous *in vitro* studies have indicated that resting M0 and polarized M2 BMDMs exhibited similar sphingolipid profiles; however, ceramide levels were predominantly elevated in M1 macrophages^40, 43^. Furthermore, AC expression was shown to decrease in M0 macrophages stimulated with LPS^43^, whereas M2 polarized macrophages had increased AC expression^40^. Though we did not measure AC expression in our studies, our assessment of AC^fl/fl^ BMDMs demonstrated similar increases in C16 ceramides. However, the sphingolipid profiles in our AC deficient M0 BMDMs were most similar to M1 macrophages, not M2; though, this may be dependent on duration of LPS/cytokine exposure. Loss of additional sphingolipid enzymes, specifically SPTLC2, has been shown to impair M1 polarized macrophages via a reduction in sphinganine levels resulting in decreased TLR4 and MyD88 association^43^. Similarly, inhibition of SPT with myriocin impaired M2 macrophage polarization and IL10 secretion by perturbing mTOR signaling^44^. However, this study did not measure sphingolipids. Lipid metabolism in a broader sense has been implicated in mediating M1/M2 macrophage polarization, in which each phenotype exhibits specific alterations in lipid composition and subsequent signaling cascades^45, 46^. For example, glycolysis and lipogenesis were upregulated in M1 macrophages increasing membrane associated lipids (including sphingolipids), whereas M2 macrophages utilized oxidative phosphorylation and fatty acid oxidation^46^. While there is an established relationship between sphingolipid metabolism and macrophage polarization, further research is needed to better understand which enzymes and metabolites drive M1/M2-like phenotypes.

Despite similarities between sphingolipid levels in M0 and M1 macrophages, we found significant discrepancies in macrophage function based on polarization state. Migration and phagocytosis decreased in M0 macrophages lacking AC, while M1 macrophages remained functional despite a reduction in polarization frequency. Interestingly, loss of AC enhanced phagocytosis in M1 macrophages. Conversely, loss of AC did not alter M2 polarization nor downstream functions such as migration or phagocytosis. Broader studies assessing the impact of lipids in macrophages have shown that manipulation of lipid regulating genes, such as SREBP or FAS, can diminish M1 proinflammatory responses by impairing lipid remodeling at the plasma membrane^46^. The M1 phenotype has been shown to be impacted by lipid remodeling while M2 macrophages are less dependent. Collectively, these suggest that AC and sphingolipid levels may differentially regulate macrophages dependent on polarization state.

Previous studies investigating the role of sphingolipids in immune cells in colitis have focused on T cells. Loss of CerS4 in T cells exacerbated weight loss in a DSS model of colitis, and increased tumorgenicity with AOM/DSS ^21^. T cell specific loss of GCS increased disease activity and histological scores, as well as colon shortening in DSS colitis^24^. Comparatively, adoptive transfer of T cells lacking either CerS6^22^ or SPTLC1^23^ protected against weight loss and histology. These data suggest that *de novo* generation of ceramides in T cells, predominantly C14 and C16 ceramides, can promote inflammation in colitis, whereas glucosylceramides may serve protective roles. We previously demonstrated that loss of AC in myeloid cells protected from weight loss in DSS colitis and reduced tumorgenicity in AOM/DSS induced CAC^12^. Although we did not measure differences in body weight or colon histology in the present study, myeloid cell-specific loss of AC protected from splenomegaly and colon shortening. These data suggest that AC in myeloid cells exhibits roles in modulating the microenvironment in colon tissues influencing both lipid metabolism and inflammation.

Changes in colonic sphingolipids have been implicated in numerous studies by our group and others in colitis and colon cancer models. T cell specific loss of CerS4 resulted in elevated C16 ceramide and HexCer in colon tissues following DSS treatment^21^. Similarly, our current study identified that loss of AC in myeloid cells elevated ceramides and SMs in colon tissues in chronic colitis. In agreement with El-Hindi et al^21^, we also identified elevated C16 HexCers, whereas sphingoid bases and S1P were reduced. Together these studies suggest the potential that sphingolipid enzymes in immune cells may influence sphingolipid metabolism in whole tissues and/or organ systems.

Our previous study identified that loss of AC in myeloid cells reduced the recruitment of neutrophils but not macrophages to the colons of mice treated with DSS^12^. The present study recapitulated these findings, and our expanded FACS panel specifically identified reductions in M1-like macrophages in the colon. Previous studies assessing loss of sphingolipid enzymes in colitis have shown that T cell-specific loss of CerS4 impaired neutrophil and macrophage recruitment into colon tissues in DSS-induced colitis; however, transfer of CerS4 deficient splenocytes elevated macrophage populations^21^. Surprisingly, T cell specific loss of CerS6 did not alter the recruitment of either Th1 or Th17 T cells as the polarization of these cell types were not altered *in vitro*^22^, although no other immune cell types were assessed. In our study, Th1 and Th17 effector T cells were reduced in the colonic tissues of AC^ΔMYE^IL10^−/-^ mice. Interestingly, *in vitro* studies have shown that neutrophils may play additional roles in driving Th1 and Th17 differentiation^47^, and in our study the reduction in effector T cells may be due in part to reduced cytokine and chemokine expression *in vivo*, as well as reduced neutrophil recruitment. Recent studies have shown that T_regs_ can acquire expression of IL17A, and that Th17 T cells may be able to trans-differentiate towards this T_regs_ phenotype as well^48, 49^. The role of IL17A+FoxP3+ T_regs_ may indicate retention of anti-inflammatory programing; however, it is challenging to discern the role of these cells in our model given the loss of IL10 production. Conversely, Th17 T cells have also shown the ability to trans-differentiate into Th1 T cells, contributing to development of colitis^50^. Indeed, our findings suggest that myeloid cell AC may exhibit an indirect role in the recruitment or maturation of effector Th1 and Th17 T cells. The collective small body of literature in conjunction with our work suggests that sphingolipid metabolism in immune cells may play a role in the mucosal recruitment of immune cells and highlights the need for further research.

Sphingolipid metabolism, specifically S1P, has been shown to regulate pSTAT3 expression in colitis. Mice treated with DSS exhibited blunted STAT3 phosphorylation after loss of either SK1^19^ or SMS2^20^, whereas loss of SK2^51^ amplified phosphorylation. Our immunohistochemistry findings indicate that loss of AC may decrease inflammation in colitis in part by reducing phosphorylation of pSTAT3. Although S1P was not significantly altered at 24 weeks of age, the partial reduction may be sufficient to reduce pSTAT3 expression given our histology data. As a known downstream target of S1P signaling, several studies have illustrated how S1P levels upregulate pSTAT3^51–54^. These studies also highlight the potential of therapeutically targeting the S1P-cytokine-pSTAT3 axis in IBD using S1P receptor modulators^55^. Furthermore, although Th1 cells have been shown to promote M1 macrophage polarization via pSTAT3 signaling^56^, these cells were reduced at 8 weeks of age in AC^ΔMYE^ mice. It may also be possible that pSTAT3 expression may be ablated due to the reduced recruitment of Th1 T cells and M1 macrophages. Collectively, our data indicate that AC may play a role in modulating STAT3-dependent pathways.

In summary, this study highlights novel cell specific roles for AC in neutrophils and macrophages. Additionally, this study indicates that loss of AC in myeloid cells is protective in chronic IBD by reducing colon tissue inflammation and impairing the recruitment of effector immune cells, suggesting AC as a potential novel therapeutic target.

**The authors have no conflicts of interest to declare**

## Author Contributions

KSE wrote the manuscript, performed conceptual and experimental design, generation of data, analysis for experiments and wrote the manuscript. BKD, JNW, and MTM: performed conceptual and experimental design, and generation of data. EMR stained, imaged, and analyzed all histological data. MRG and CLD: generated data and analysis of experiments. CAB and PDD: contributed to the experimental design. JMS, RJS, JEW, and PRK: assisted with data analysis and manuscript editing. AJS conceived the original hypothesis, provided necessary funding for the materials and methods, and supervised the project.

## Acknowledgements

Histology utilized in this publication was provided by the UACC TACMASR and lipidomic analyses were provided by the University of Arizona Cancer Center Analytical Chemistry Shared Resource Core both supported by the National Cancer Institute P30 CA023074. Flow cytometry data in this publication were obtained using the BD LSR Fortessa flow cytometer provided by the University of Arizona Department of Pediatrics.

