## Supplemental Table for "Loss of Acid Ceramidase in Myeloid Cells Protects from Chronic Colitis in IL10-Deficent Mice"

**Supplementary Table 1**

| **Item** | **Vendor** | **Catalog#** | **Dilution** |
| --- | --- | --- | --- |
| **RNA AND PCR MATERIALS** | | | |
| PureLink RNA isolation kit | Thermo | 12183025 | NA |
| qScript cDNA Synthesis Kit | VWR | 101414-106 | NA |
| Mouse CCL2 primer - FAM | Thermo | Mm00441242_m1 | NA |
| Mouse CXCL2 primer - FAM | Thermo | Mm00436450_m1 | NA |
| Mouse CCL5 primer - FAM | Thermo | Mm01302427_m1 | NA |
| Mouse S100a9 primer - FAM | Thermo | Mm00656925_m1 | NA |
| Mouse IL-1β primer - FAM | Thermo | Mm00434228_m1 | NA |
| Mouse CXCL1 primer - FAM | Thermo | Mm04207460_m1 | NA |
| Mouse MMP8 primer - FAM | Thermo | Mm00439509_m1 | NA |
| Mouse MMP9 primer - FAM | Thermo | Mm00442991_m1 | NA |
| Mouse F4/80 primer - FAM | Thermo | Mm00802529_m1 | NA |
| Mouse TNFα primer - FAM | Thermo | Mm00443260_g1 | NA |
| Mouse OCLN primer - FAM | Thermo | Mm00500912_m1 | NA |
| Mouse CDH1 primer - FAM | Thermo | Mm01247357_m1 | NA |
| Mouse TJP1 primer - FAM | Thermo | Mm00493699_m1 | NA |
| Mouse COX2 primer - FAM | Thermo | Mm00478374_m1 | NA |
| Mouse Beta Actin primer - VIC | Thermo | Mm00607939_s1 | NA |
| **ELISA MATERIALS** | | | |
| EDTA-free protease inhibitor | Millipore Sigma | 11836170001 | NA |
| ELISA buffer kit | Thermo | 900K00 | NA |
| ELISA Maxisorp Plates | Thermo | 439454 | NA |
| IL-6 Mouse ELISA | Thermo | 900-K50K | NA |
| CCL2 Mouse ELISA | Thermo | 900-K126K | NA |
| CXCL2 Mouse ELISA | Thermo | 900-K152K | NA |
| TNFα Mouse ELISA | Thermo | 900K54 | NA |
| IFNg Mouse ELISA | Thermo | 900-K98 | NA |
| IL-1b Mouse ELISA | Thermo | 900-K47 | NA |
| **IMMUNOHISTOCHEMISTRY MATERIALS** | | | |
| pSTAT3 (Tyr705, D3A7) rabbit anti-mouse monoclonal antibody | Cell Signaling Technology | 9145 | 1:250 |
| Ki67 (SP6) rabbit anti-mouse monoclonal antibody | Fisher Scientific | NC1494892 | 1:500 |
| COX2 (EPR12012) rabbit anti-mouse monoclonal antibody | abcam | Ab179800 | 1:3000 |
| Mayer’s Hematoxylin | Fisher | NC1758756 | NA |
| Rabbit Specific IHC polymer detection kit HRP/DAB | Fisher | NC1924340 | NA |
| Pierce DAB Substrate Kit | Thermo | 34002 | NA |
| **FLOW CYTOMETRY MATERIALS** | | | |
| RBC Lysis Buffer | Fisher | 420301 | NA |
| Collagenase type 1 | Fisher | 17018-029 | NA |
| FC Block | Fisher | NC1926472 | 1:50 |
| Live/Dead Fixable Stain DAPI | Fisher | L34961 | 1:4000 |
| CD45-BV510 rat IgG2a κ | Fisher | BDB56389 | 1:100 |
| CD3-BUV496 rat IgG2b κ | Fisher | BDB741117 | 1:200 |
| CD4-PE-Cy7 rat IgG2a κ | Fisher | BDB552775 | 1:400 |
| CD8-BV750 rat IgG2a κ | Fisher | BDB747134 | 1:200 |
| CD19-PerCP-Cy5.5 rat IgG2a κ | Fisher | BDB551001 | 1:100 |
| CD11b-FITC rat IgG2b κ | Fisher | 509436 | 1:100 |
| F4/80-APC, rat IgG2a κ | Fisher | 50-112-8925 | 1:50 |
| MHCII-PacificBlue rat IgG2b k | Fisher | 50-163-190 | 1:100 |
| CD11c-APC-eF780 Armenian Hamster IgG | Fisher | 50-161-43 | 1:100 |
| LY6-G-PE rat IgG2a κ | Fisher | BDB551461 | 1:200 |
| Ly6C-PE-Cy7 rat IgM k | Fisher | BDB560593 | 1:100 |
| CD206-PE-Cy5 rat IgG2a k | Fisher | 50-238-8273 | 1:100 |
| Permeabilization buffer | Fisher | 501129059 | NA |
| CD25 APC rat IgG1 λ | Fisher | BDB561048 | 1:100 |
| FoxP3 PE Rat IgG2b k | Fisher | 50-164-290 | 1:250 |
| IL-17 AF488 rat IgG1 k | Fisher | 50-170-001 | 1:250 |
| IFNγ PacBlue rat IgG1 | Fisher | 50-169-929 | 1:250 |
| GranzymeB APCeF780 rat IgG2a k | Fisher | 47-889-880 | 1:250 |
| Round Bottom FACS tubes | Fisher | 352054 | NA |
| 96 well round bottom plate | Sarstedt | 82.1582.001 | NA |
| FlowJo v10.8 | BD Life Sciences | NA | NA |
| **CELL CULTURE MATERIALS** | | | |
| 10cm non-cell culture treated petri dishes | VWR | 25373-100 | NA |
| Mouse M-CSF | Thermo | 315-02-50UG | NA |
| EasySep™ Mouse Neutrophil Enrichment Kit | Stem Cell | 19762 | NA |
| LPS | Millipore Sigma | L4391-1MG | NA |
| Calcein AM | VWR | 89044-504 | NA |
| Corning BioCoat FluoroBlok 3μm 24 well transwell inserts | VWR | BD354597 | NA |
| Mouse C5a | Thermo | 315-40-5UG | NA |
| Mouse MCP1 | Thermo | 250-10-10UG | NA |
| Phagocytosis assay kit IgG FITC | Cayman Chemical | 500290 | NA |
| Mouse IFNγ | Thermo | 315-05-20UG | NA |
| Mouse IL-4 | Fisher | 50-170-363 | NA |
| Mouse IL-13 | Fisher | 50-170-418 | NA |
| **OTHER MATERIALS** | | | |
| FITC-Dextran | Millipore Sigma | 68059-1G | NA |
| GraphPad Prism v10.6 | GraphPad | NA | NA |
