## Supplemental Figures for "Loss of Acid Ceramidase in Myeloid Cells Protects from Chronic Colitis in IL10-Deficent Mice"

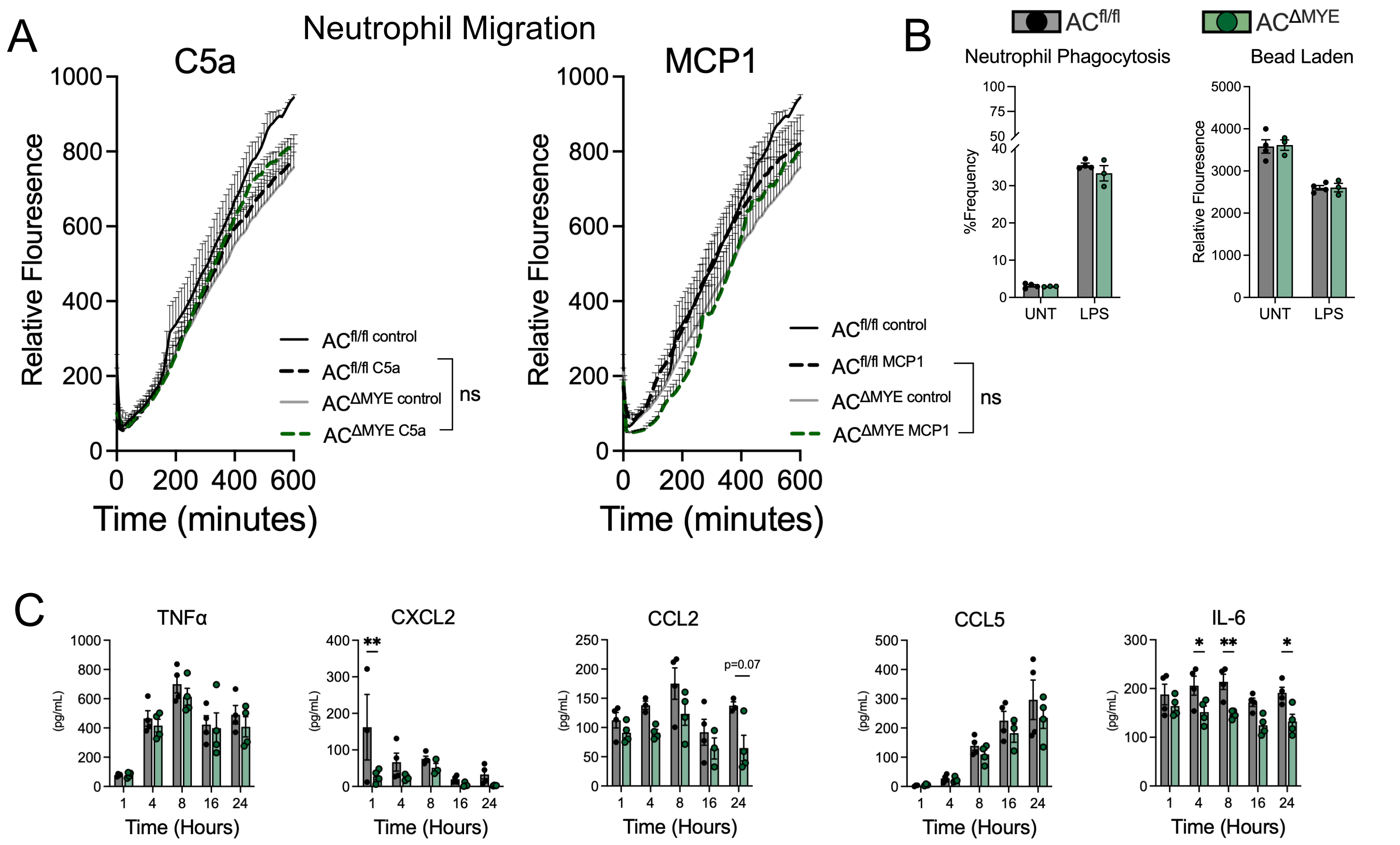


Supplemental Figure 1. AC does not regulate inflammatory functions in neutrophils. (A) BMDN migration assay, terminal migration timepoint assessed by 2-way ANOVA. (B) BMDN phagocytosis assay, 2-way ANOVA, Fisher's LSD test; phagocytic cells (left) and bead laden quantification (right). (C) Post-LPS stimulation cytokine and chemokine media assessment via ELISA, 2-way ANOVA, Šídák multiple comparisons correction. Data represent mean ± SEM, n=4 ; *p<0.05, **p<0.01, ***p<0.001.

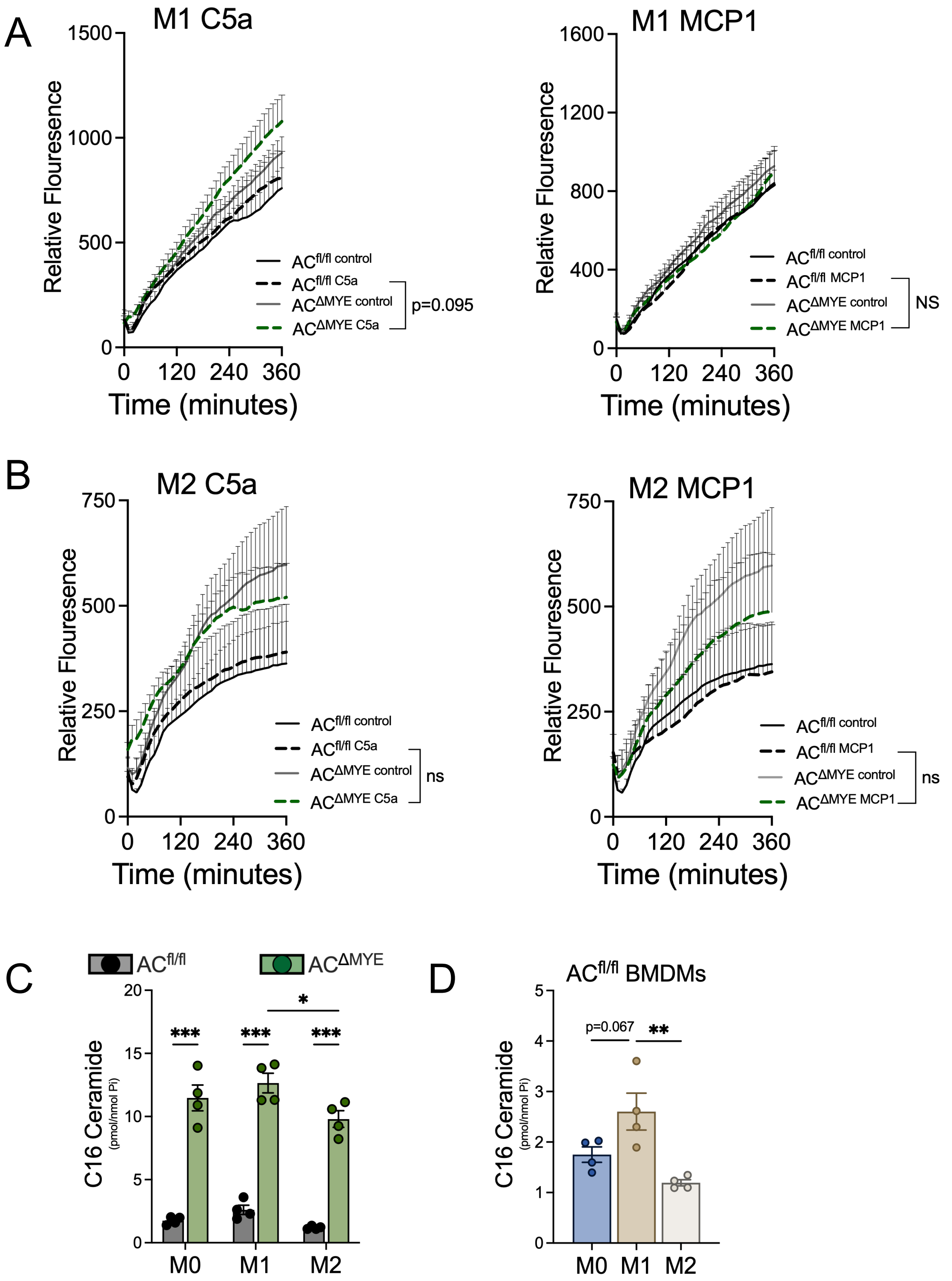


Supplemental Figure 2. AC does not regulate polarized cell migration. Migration assays in (A) M1 and (B) M2 polarized BMDMs, terminal migration timepoint assessed by 2-way ANOVA. (C) C16 ceramide comparisons between genotypes and polarization state, 2-way ANOVA. (D) C16 ceramide abundance based on polarization state in $AC^{fl/fl}$ BMDMs, 1-way ANOVA. Data represent mean ± SEM, n=3-4; *p<0.05, **p<0.01, ***p<0.001.

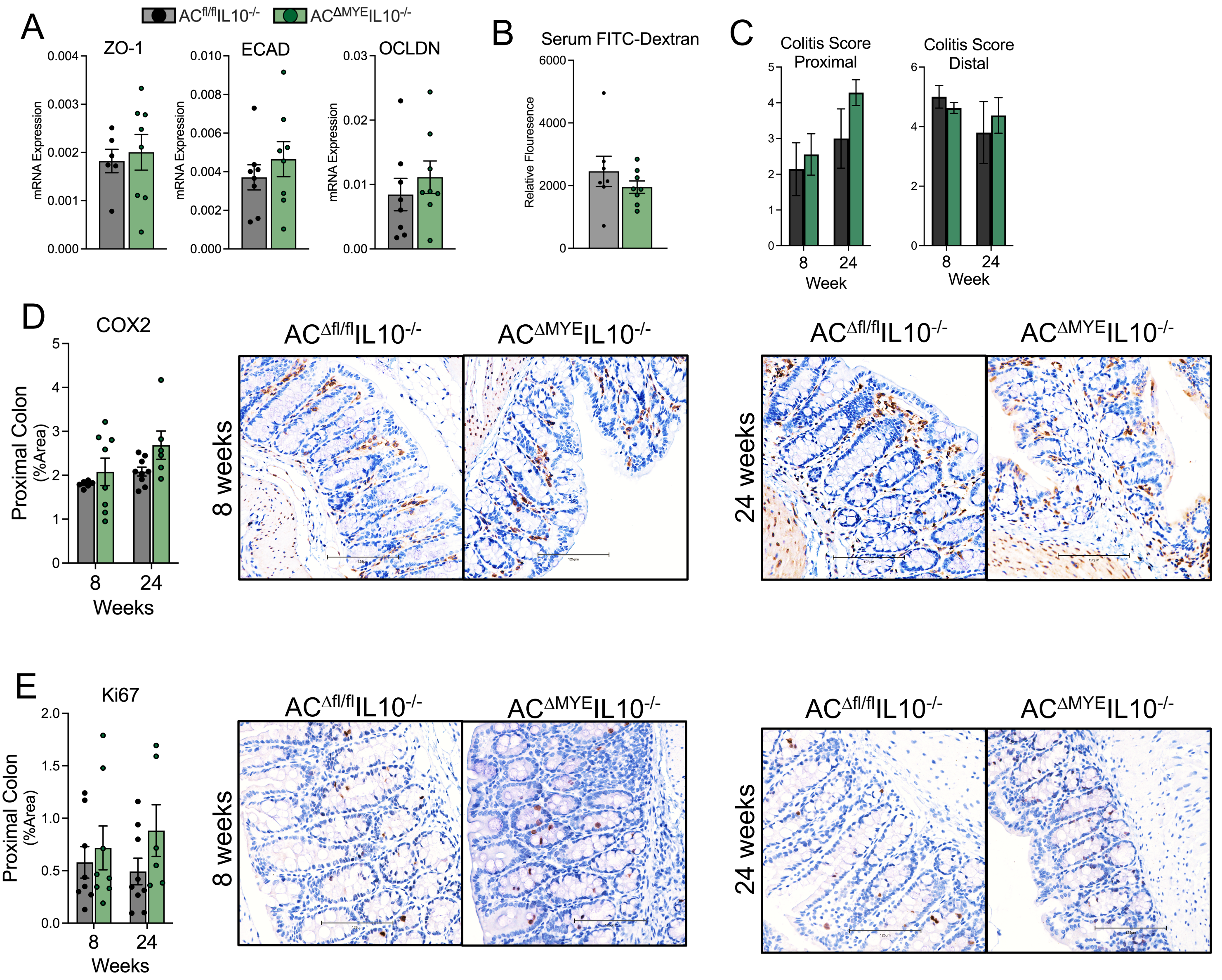


Supplemental Figure 3. AC does not alter intestinal permeability or histology. (A) rt-PCR mRNA analysis of colon tissue tight junction protein expression at 24 weeks of age, n≥7, unpaired t test. (B) Serum FITC dextran at 24 weeks of age, n≥7, unpaired t test. (C) Colitis severity scoring from H&E colon tissue (histology not shown). Histological assessment of proximal colon tissue expression of (D) COX2 and (E) Ki67, n≥6. Representative images taken at 20x. Data represent mean ± SEM; *p<0.05, **p<0.01, ***p<0.001.

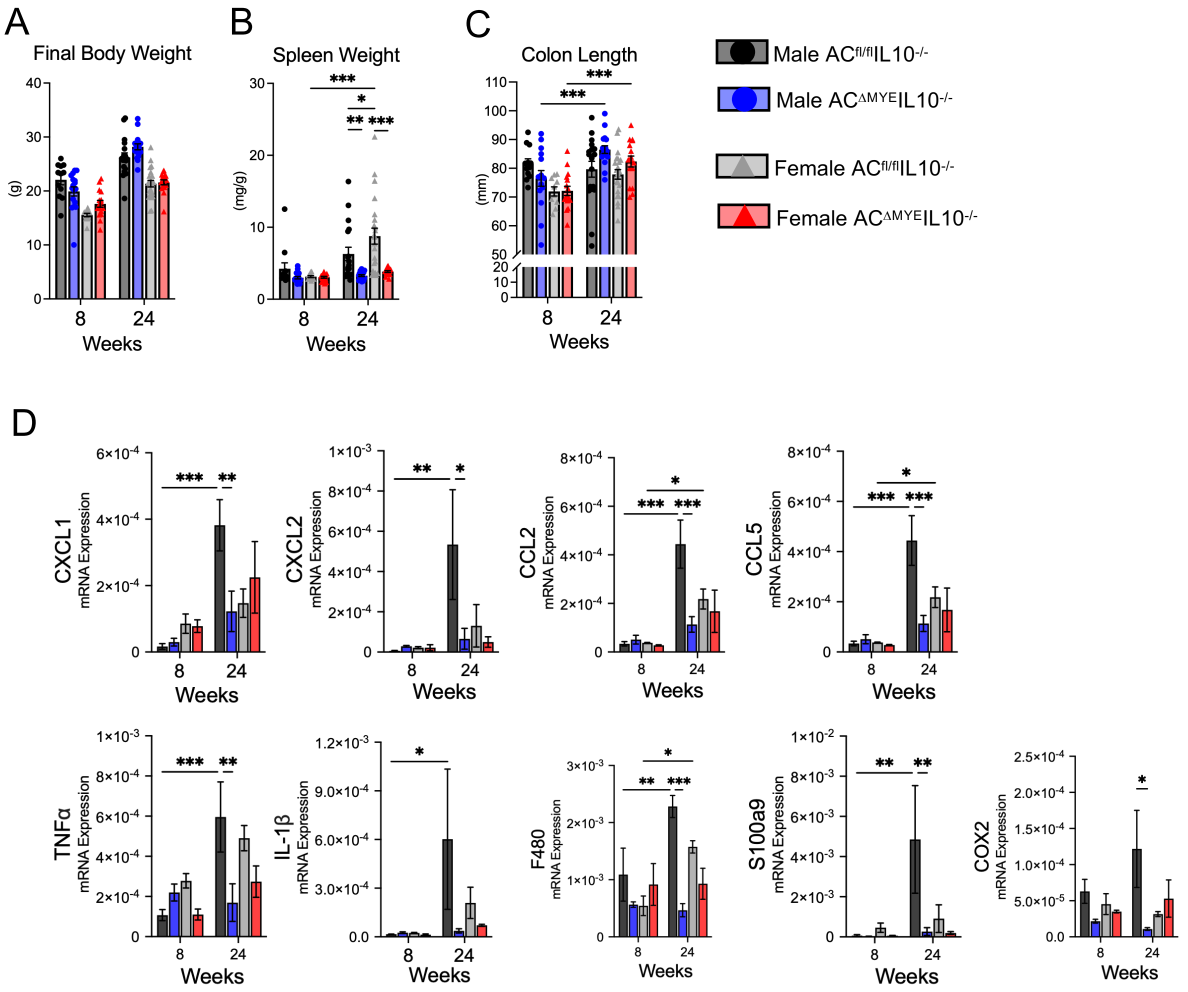


Supplemental Figure 4. Protection from chronic colitis exhibits sex differences. (A) Terminal body weight, (B) spleen weight normalized to body weight, and (C) colon length, n≥11. (D) XBP1 splicing quantification, n≥4. (E) rt-PCR mRNA analysis of colon tissue, n≥3. 2-way ANOVA with Tukey multiple comparisons correction. Data represent mean ± SEM; *$p<0.05$, **$p<0.01$, ***$p<0.001$.

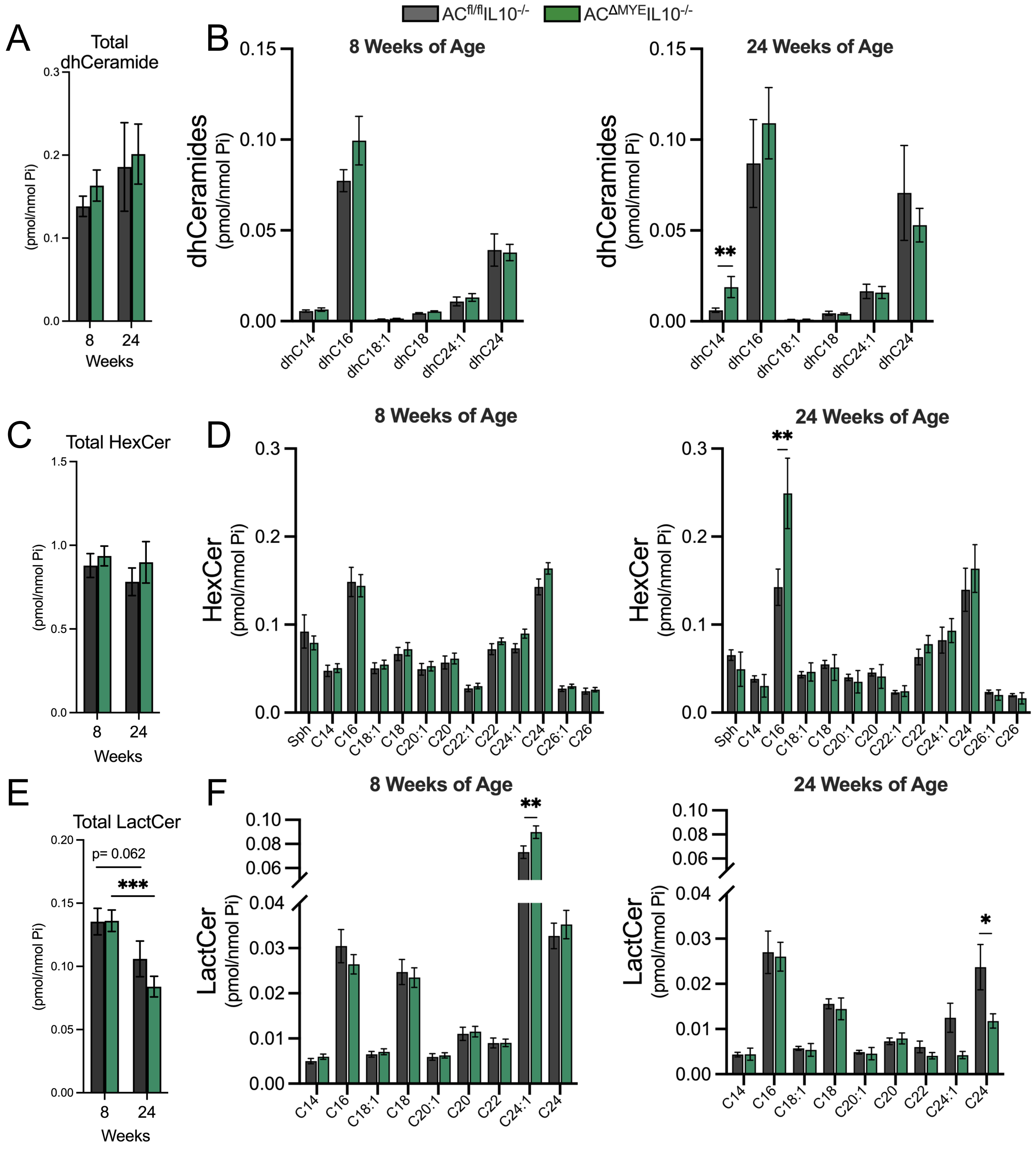


Supplemental Figure 5. AC does not significantly regulate colon tissue de novo sphingolipids or glucosylceramides (GluCer). (A) Total dhceramides and (B) species distribution at 8 or 24 weeks of age. (C) Total hexosylceramides (HexCer) and (D) species distribution at 8 or 24 weeks of age. (E) Total lactosylceramides (LactCer) and (F) species distribution at 8 or 24 weeks of age. 2-way ANOVA, Fisher's LSD test. Data represent mean ± SEM, n≥8; *p<0.05, **p<0.01, ***p<0.001.

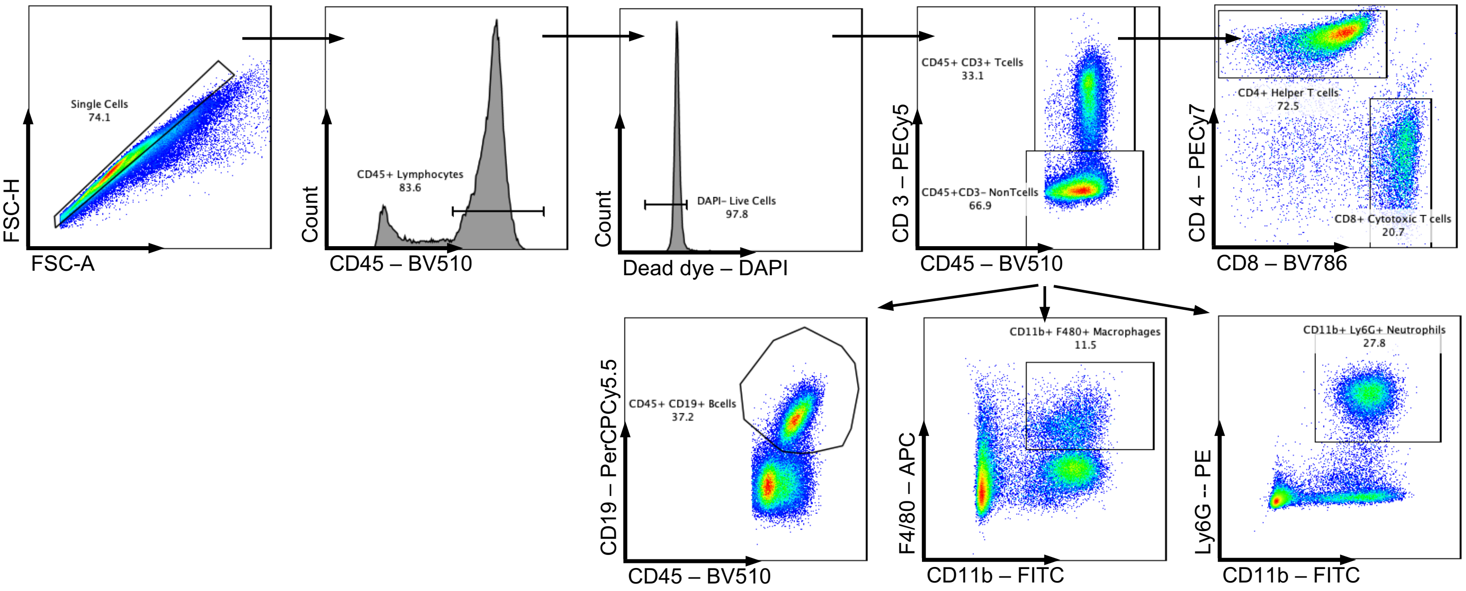


Supplemental Figure 6. Flow cytometry gating schemes for basic immunophenotyping in main figure 5 and supplemental figure 7.

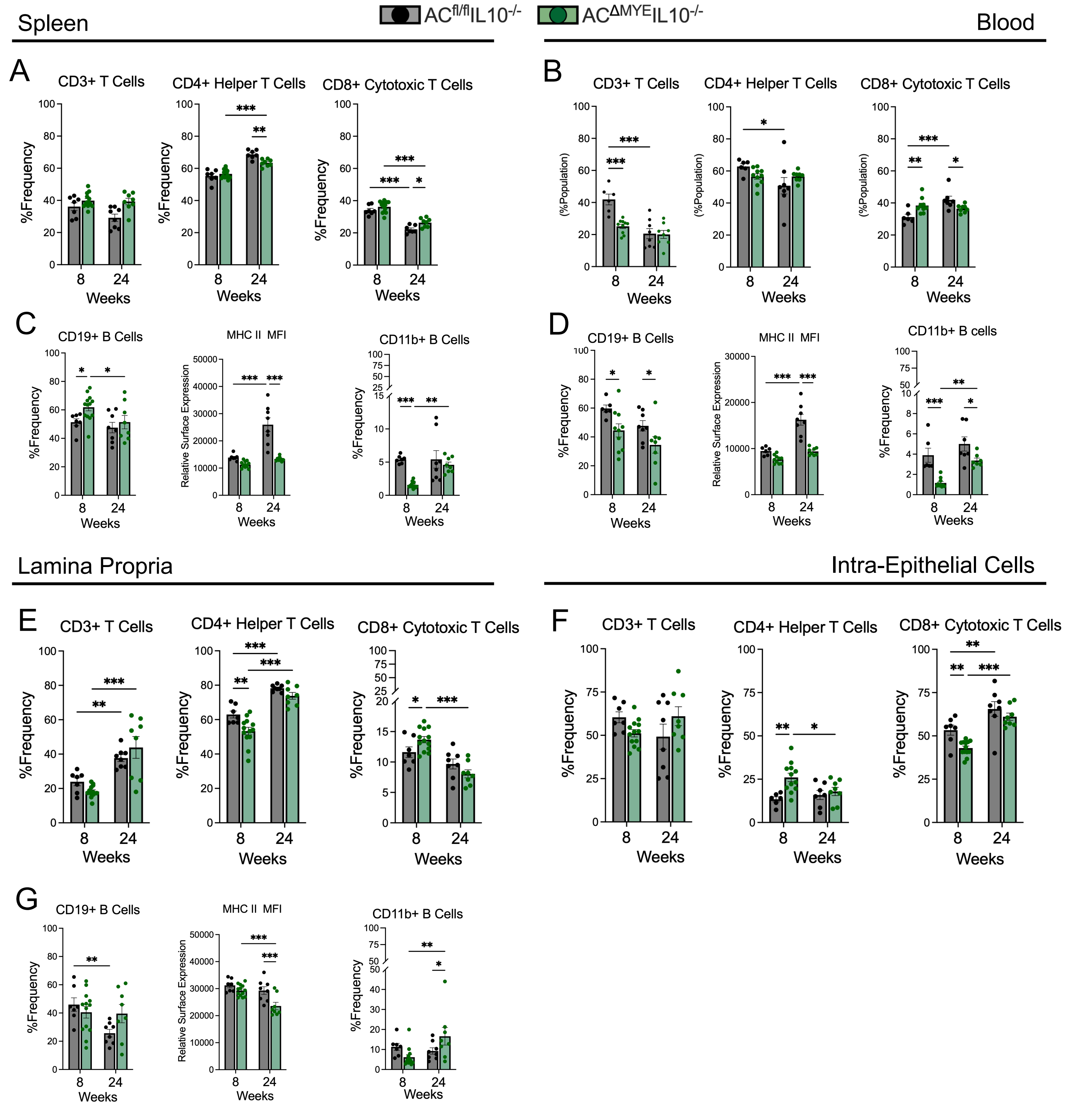


Supplemental Figure 7. Myeloid cell AC alters lymphocyte populations. T cell phenotyping in (A) spleen, (B) blood, and (E) LP and (F) IEC colon tissue compartments. B cell phenotyping in (C) spleen, (D) blood, and (G) colonic LP. 2-way ANOVA, Fisher's LSD test. Data represent mean ± SEM, n≥6; *p<0.05, **p<0.01, ***p<0.001.

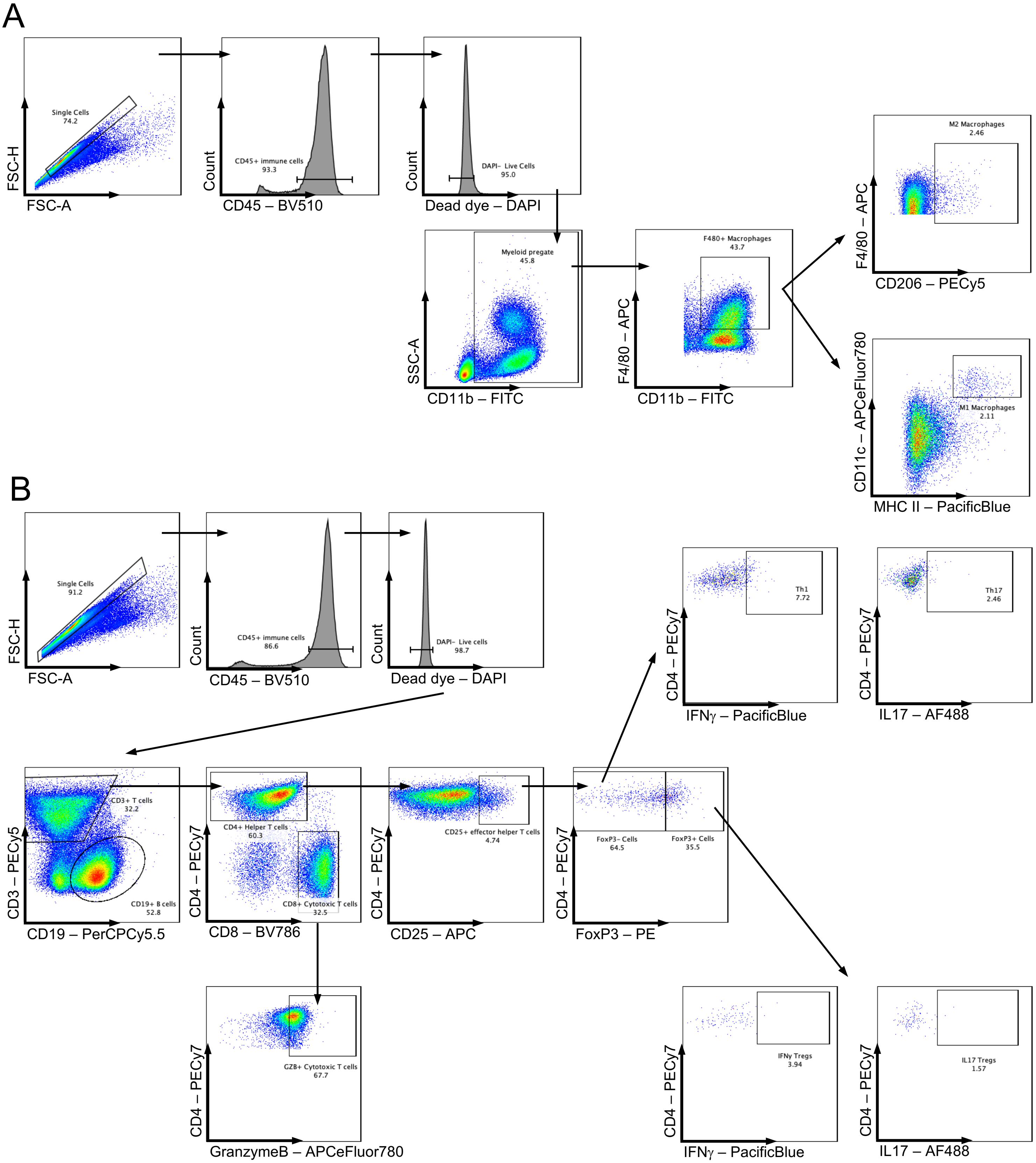


Supplemental Figure 8. Gating schemes for effector cell phenotyping. (A) Gating scheme for M1/M2 macrophages and (B) effector T cells in main figure 6 and supplemental figure 9.

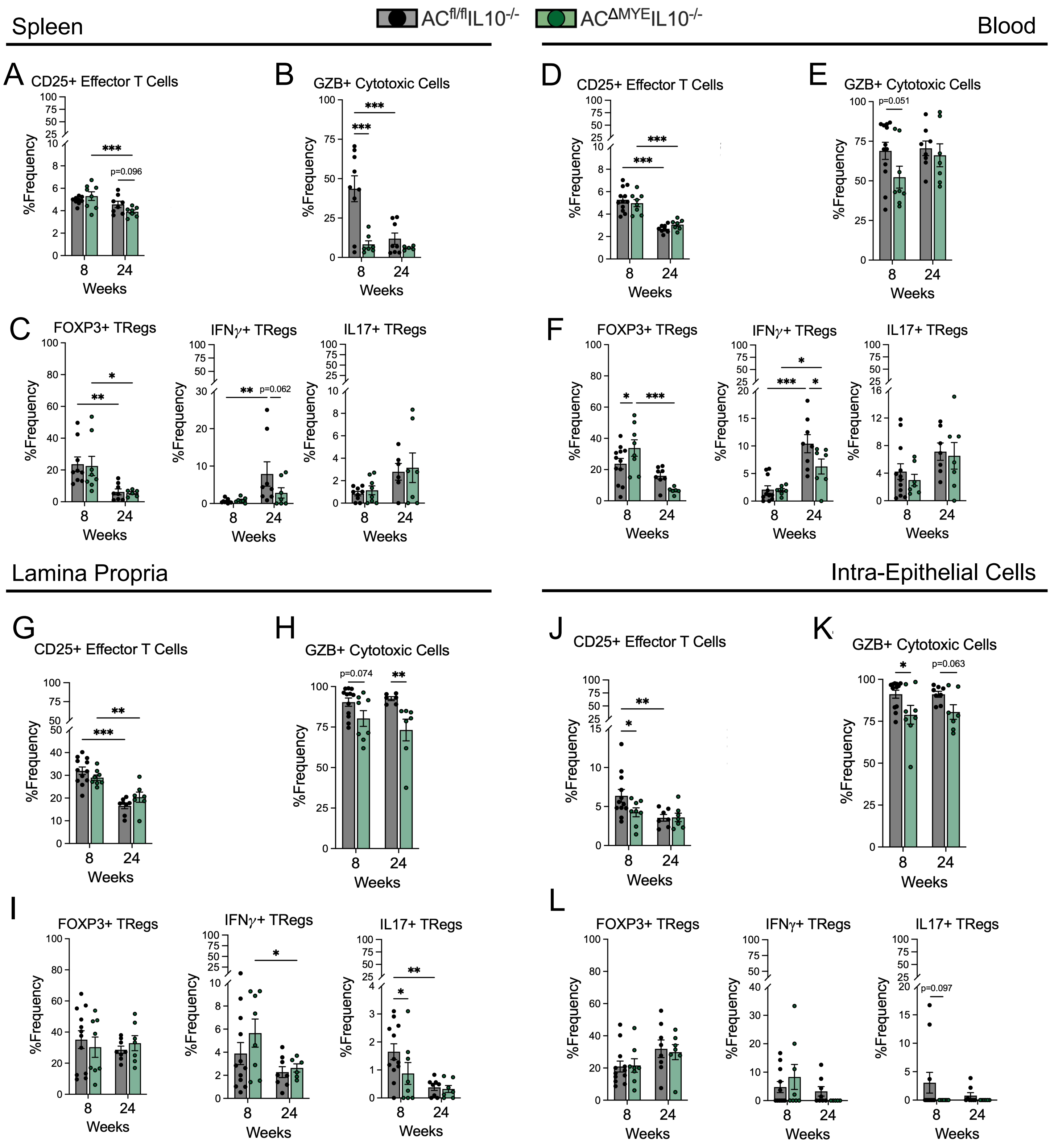


Supplemental Figure 9. Myeloid cell AC regulates effector and regulatory T cells throughout tissues. CD4+CD25+ effector T cells in (A) spleen, (D) blood, and (G) LP and (J) IEC colon tissue compartments. CD8+GranzymeB+ cytotoxic T cells in (B) spleen, (E) blood, and (H) LP and (K) IEC colon tissue compartments. CD4+CD25+FoxP3+ regulatory T cell phenotyping and subtyping in (C) spleen, (F) blood, and (I) LP and (L) IEC colon tissue compartments. 2-way ANOVA, Fisher's LSD test. Data represent mean ± SEM, n≥6; *$p<0.05$, **$p<0.01$, ***$p<0.001$.
